# Morphogenetic Patterning During Regional and Cell Type Specification in the Embryonic Basal Ganglia

**DOI:** 10.64898/2026.04.14.718571

**Authors:** Jia Sheng Hu, Karol Cichewicz, Jonathan W. C. Lim, Linda J. Richards, Luis Puelles, Alex S. Nord, John L. Rubenstein

## Abstract

Cell type specification in the embryonic brain and spinal cord is thought to begin within molecularly defined progenitor domains that do not intermix. Our data provide an alternative model that is spatially and temporally dynamic within a basal ganglia anlage, the medial ganglionic eminence (MGE). MGE progenitor cells are progressively displaced ventrally and caudally from a rostral growth zone (the MGE/LGE sulcus). Progenitors that leave the MGE/LGE sulcus early occupy caudoventral MGE regions, while ones that leave later reside in rostrodorsal MGE regions. As they change position, their transcriptional states and cell type output change. Transcriptional analyses showed an upregulation of the *Nfi* TFs during the period of progenitor movement. *Nfia* and *Nfib* double mutants alter the repertoire of cortical interneuron subtypes. Overall, we present a mechanism that synchronizes regional patterning with tissue growth and links spatial and temporal specification in producing diverse neuronal subtypes.

## Introduction

Prevailing evidence proposes that cell type specification in the embryonic brain and spinal cord begins within molecularly defined progenitor domains (ventricular and subventricular zones (VZ, SVZ)) that do not intermix (*1-3*). Within the basal ganglia anlagen, the medial ganglionic eminence (MGE) has distinct domains of transcription factor (TF) expression that are thought to participate in the specification of subtypes of pallidal projection neurons (PN) [e.g. globus pallidus (GP) and some BST components] and interneurons (IN) that migrate to the striatum and cortex (*1*,*4*).

Our data provide an alternative model that is spatially and temporally dynamic. During early stages, MGE VZ cells are progressively displaced ventrally and caudally from a rostral growth zone located at the MGE/LGE sulcus. As they move, their transcriptional states and cell type output change. Transcriptional analyses show temporal changes as the progenitors move, including the induction of the *Nfi* TFs. *Nfia* and *Nfib* double mutants (dcKO) alter the repertoire of IN subtypes. We suggest that neuroembryology needs to re-evaluate how tissue growth and cell type specification are coordinated over developmental time.

## Results

### Morphogenetic regional-temporal patterning of the MGE VZ: Progenitors from the rostrodorsal MGE move caudoventraly along the VZ

To study the properties and fates of progenitors from the MGE/LGE sulcus, we used the stable transgenic enhancer line, *1538*, that drives the expression of a tamoxifen-inducible *Cre* (*CreERT2*) and *GFP* (*5*). *GFP* expression was used to detect ongoing *1538* enhancer activity, while *CreERT2* was used to fate map *1538* lineages from various ages. *1538* is probably a *Nkx2*.*1* enhancer (*5*), which has VZ activity in two subpallial domains: the MGE/LGE sulcus (green arrow) (contiguous with the pallidal paraseptal region (PSe)), and the preoptic area (POA) from E9.5 thru E13.5 (Fig. 1A and fig. S1).

**Fig. 1.**
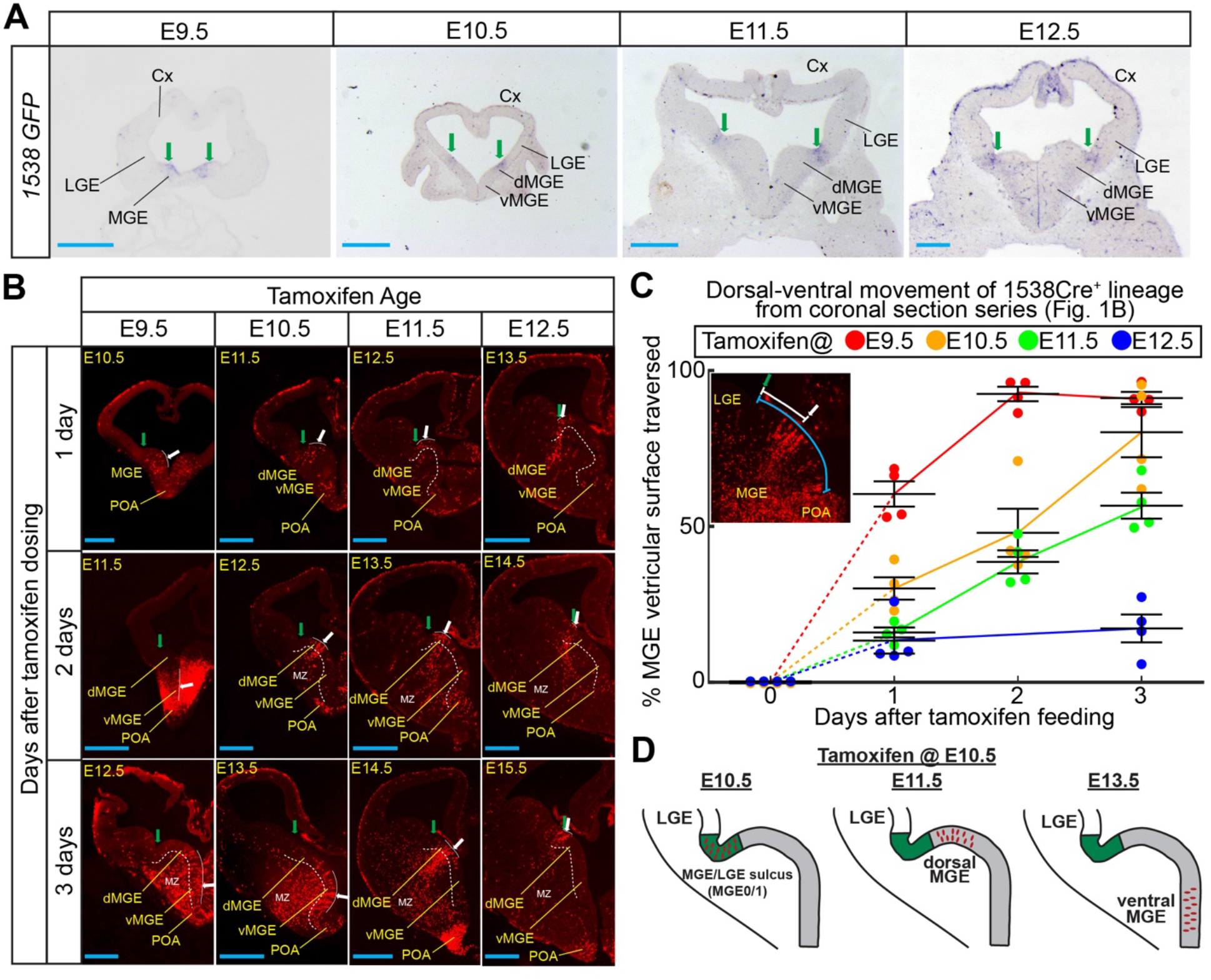
MGE progenitors in the 1538Cre-lineage move ventrally in the VZ. **(A)** *GFP in situ* hybridization (ISH) on E9.5 thru E12.5 coronal sections reporting 1538 activity. *GFP* expression remains in the MGE/LGE sulcus (green arrow; also known as MGE0/1 (*2*)) at these ages. Blue scale bar, 200 uM. **(B)** Fate mapping of 1538 lineages (tdTomato^+^) one, two, and three days after tamoxifen treatment. Progenitors of the E9.5, E10.5, and E11.5 1538 lineages (white arrow) move ventrally away from the MGE/LGE sulcus (green arrow) along the MGE VZ. White dash line, MGE SVZ and MZ boundary. White solid line, distribution of 1538 lineages in the VZ. Blue scale bar, 200 uM. **(C)** Measurement of E9.5 (red), E10.5 (orange), E11.5 (green), and E12.5 (blue) 1538 progenitor movement along the ventricular surface 1, 2, and 3 days after tamoxifen treatment (n=4 for each time point). Inset: 1538 movement was calculated by measuring the distance (white line) between the center of the 1538 progenitors (white arrow) and the MGE/LGE sulcus (green arrow) and normalizing it to the distance (blue line) between the MGE/LGE and MGE/POA sulcus. Error bars, SEM. **(D)** Illustration of progenitors from the E10.5 1538 lineage moving from the MGE/LGE sulcus (green) into the ventral MGE (grey). Abbreviations: dMGE, dorsal medial ganglionic eminence; LGE, lateral ganglionic eminence; MGE, medial ganglionic eminence; MZ, mantle zone; POA, preoptic area; vMGE, ventral medial ganglionic eminence.

The fate of *1538* active cells from the VZ of the MGE/LGE sulcus was followed by tamoxifen administration at either E9.5, E10.5 or E11.5 to pregnant *1538*^+/-^; Ai14^f/f^ (*tdTomato* Cre reporter) (*6*) mice. We assessed the position of tdTomato^+^ cells 1, 2, and 3 days after tamoxifen administration. Surprisingly, one day after tamoxifen, *1538* progenitor lineages were no longer at the MGE/LGE sulcus but had moved ventrally (white arrow) along the MGE VZ (Fig. 1B, fig. S2); no dorsal movement into the LGE was detected. Furthermore, two and three days after tamoxifen, these labeled progenitor lineages moved even further ventrally (white arrows). However, by E12.5, VZ movement away from the MGE/LGE sulcus ceased (Fig. 1B). In contrast, no VZ movement was detected from the POA at all ages examined (Fig. 1B).

We measured the ventral movement of *1538* E9.5-E12.5 progenitors from the MGE/LGE sulcus (Fig. 1C). The E9.5 lineage moved the fastest, while E10.5 and E11.5 were intermediate, and E12.5 moved the slowest.

We used time-lapse imaging of slice cultures (from E10.5 tamoxifen-treated *1538*^+^ embryos) to investigate cell movement away from the MGE/LGE sulcus along the VZ. At E11.5, telencephalons were coronally sliced, cultured and imaged. By 18 and 24 hours *in vitro*, tdTomato^+^ cells moved along the VZ (white arrow) away from the MGE/LGE sulcus (green) (fig. S3). Taken together, progenitors generated at E9.5-E11.5 in the MGE/LGE sulcus, moved ventrally in the MGE VZ, providing evidence that the MGE VZ has lateral movement during these stages (Fig. 1D).

We then assessed whether the *1538* lineage also moved rostrocaudally along the MGE VZ. E10.5 *1538* lineages were analyzed sagittally 1 and 3 days after tamoxifen dosing. By 3 days after tamoxifen, in caudal regions, this lineage reached the dorsal border of the POA (fig. S4). Therefore, MGE VZ cells move both dorsal to ventral and rostral to caudal.

We wondered if the MGE/LGE sulcus is a growth zone with high mitotic index that drives MGE morphogenesis by generating cohorts of VZ progenitors that are displaced caudoventrally. Accordingly, we used M-phase pH3 labelling to test whether the dorsal MGE had a higher mitotic index than the ventral MGE. At E10.5 and E11.5 the mitotic index of the dorsal MGE was higher than the ventral MGE (Fig. 2A, fig. S5). Of note, at E13.5 when the 1538 VZ movement has stopped, there was no longer a higher mitotic index in the MGE/LGE sulcus (fig. S5). Thus, we postulate that the MGE/LGE sulcus is a growth zone: progenitors that arise from the sulcus displace older VZ cells caudoventrally.

**Fig. 2.**
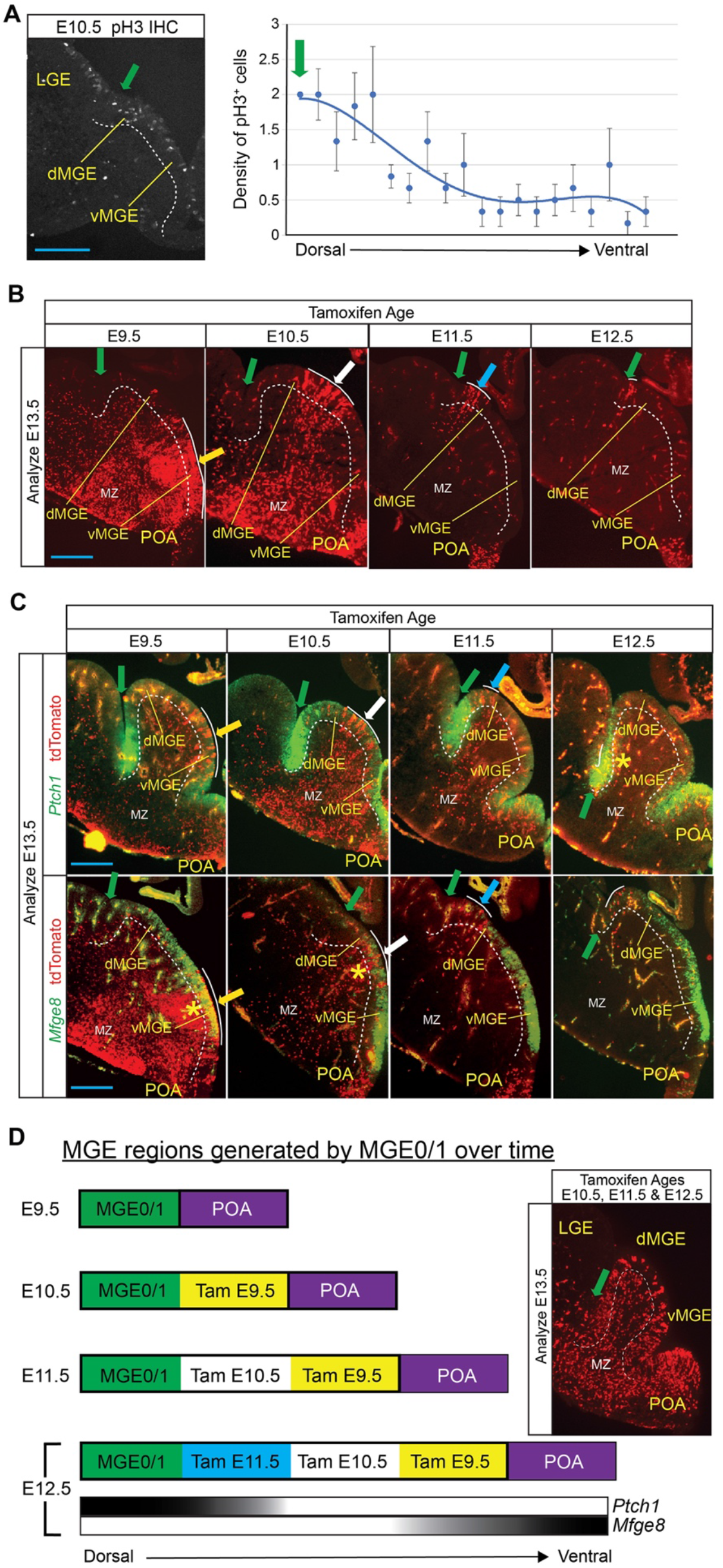
1538 lineages moving out of the MGE/LGE sulcus sequentially pattern the MGE. **(A)** phospho-histone H3 (pH3) immunohistochemistry on a E10.5 coronal wildtype section (left panel). pH3 density measurement along the MGE VZ (right panel) (n=6). Density was counted in 20 equally spaced bins along the dorsal-ventral MGE VZ axis. Green arrow: MGE/LGE sulcus. Note, higher pH3 density in the MGE/LGE sulcus. Blue scale bar, 200 uM. Error bars, SEM. **(B)** Fate mapping at E13.5 of 1538 lineages (tdTomato^+^) based on tamoxifen-dosing at different ages: E9.5, E10.5, E11.5 and E12.5. Green arrow, MGE/LGE sulcus. Yellow, white, and blue arrows indicate the center of E9.5, E10.5 and E11.5 1538 lineages, respectively. Blue scale bar, 200 uM. **(C)** *Ptch1* (top row, green) and *Mfge8* (bottom row, green) fluorescent ISH co-stained with tdTomato (red) at E13.5 comparing 1538 lineages following tamoxifen at E9.5, E10.5, E11.5 and E12.5. Asterisk, region of tdTomato and *Ptch1* or *Mfge8* coexpression. Yellow, white, and blue arrows indicate the approximate center of E9.5, E10.5 and E11.5 1538 lineages within the VZ, respectively. Blue scale bar, 200 uM. **(D)** Schema illustrating how during MGE growth the earliest 1538 lineages occupy the ventral-most positions in the MGE (*Mfge8*^+^, *Ptch1*^-^) while later 1538 lineages occupy progressively more dorsal positions (*Mfge8*^-^, *Ptch1*^+^). Fate mapping of 1538 lineages at E13.5 after giving tamoxifen at E10.5, E11.5, and E12.5 to the same animal (inset). Green arrow, MGE/LGE sulcus. White dash line, MGE SVZ and MZ boundary. White solid line, distribution of 1538 lineages in the VZ. Abbreviations: dMGE, dorsal medial ganglionic eminence; LGE, lateral ganglionic eminence; MGE, medial ganglionic eminence; MZ, mantle zone; POA, preoptic area; vMGE, ventral medial ganglionic eminence.

Next, we hypothesized that cohorts of VZ cells produced by the MGE/LGE sulcus at early times move away larger distances than cells produced at later times. Indeed, the 1538 lineage generated at E9.5 or E10.5 moved further ventrally than lineages generated at E11.5 or E12.5 (Fig. 2B). Given that different domains within the MGE VZ express distinct sets of genes (*1*,*2*), progenitors may change their transcriptomes as they move ventrally. To test this, we performed double-labeling of tdTomato with either a marker of the MGE/LGE sulcus (*Ptch1)* or of the ventral MGE VZ (*Mfge8*) at E13.5 (Fig. 2C). Whereas the early lineages (tamoxifen at E9.5, E10.5) express *Mfge8*, the latest lineage (tamoxifen at E12.5) expresses *Ptch1*. To further test this hypothesis, we gave tamoxifen at E10.5, E11.5, and E12.5 and assessed the position of tdTomato^+^ cells at E13.5 (Fig 2D). We found that tdTomato^+^ cells from successive ages populate the full dorsal-ventral extent of the MGE. Taken together, we propose a morphogenetic model whereby the E9.5-E12.5 MGE grows asymmetrically from cells largely generated in the MGE/LGE sulcus that successively populate more ventral and caudal regions (Fig. 2D).

### Progenitors originating in the rostrodorsal MGE change gene expression as they move caudoventrally along the MGE VZ

Previous studies have used single cell RNAseq to characterize MGE cell types (*7-10*). Here, we characterized the transcriptomes of MGE progenitors as they move caudoventrally away from the MGE/LGE sulcus. To this end, we gave tamoxifen to 1538 mice at E10.5 and dissected the MGE at either E11.5 or E13.5 (when tdTomato cells were in the dorsal and ventral MGE positions, respectively) (Fig. 2). TdTomato^+^ cells were purified by FACS; their RNAs were isolated and used to generate single-cell RNA-seq (scRNAseq) libraries. Their transcriptomes were integrated and projected onto the same UMAP (Fig. 3A). The distribution of *Hes1, Ascl1* and *Dcx-*expression approximates the locations of VZ, SVZ and MZ cells (Fig. 3A).

**Fig. 3.**
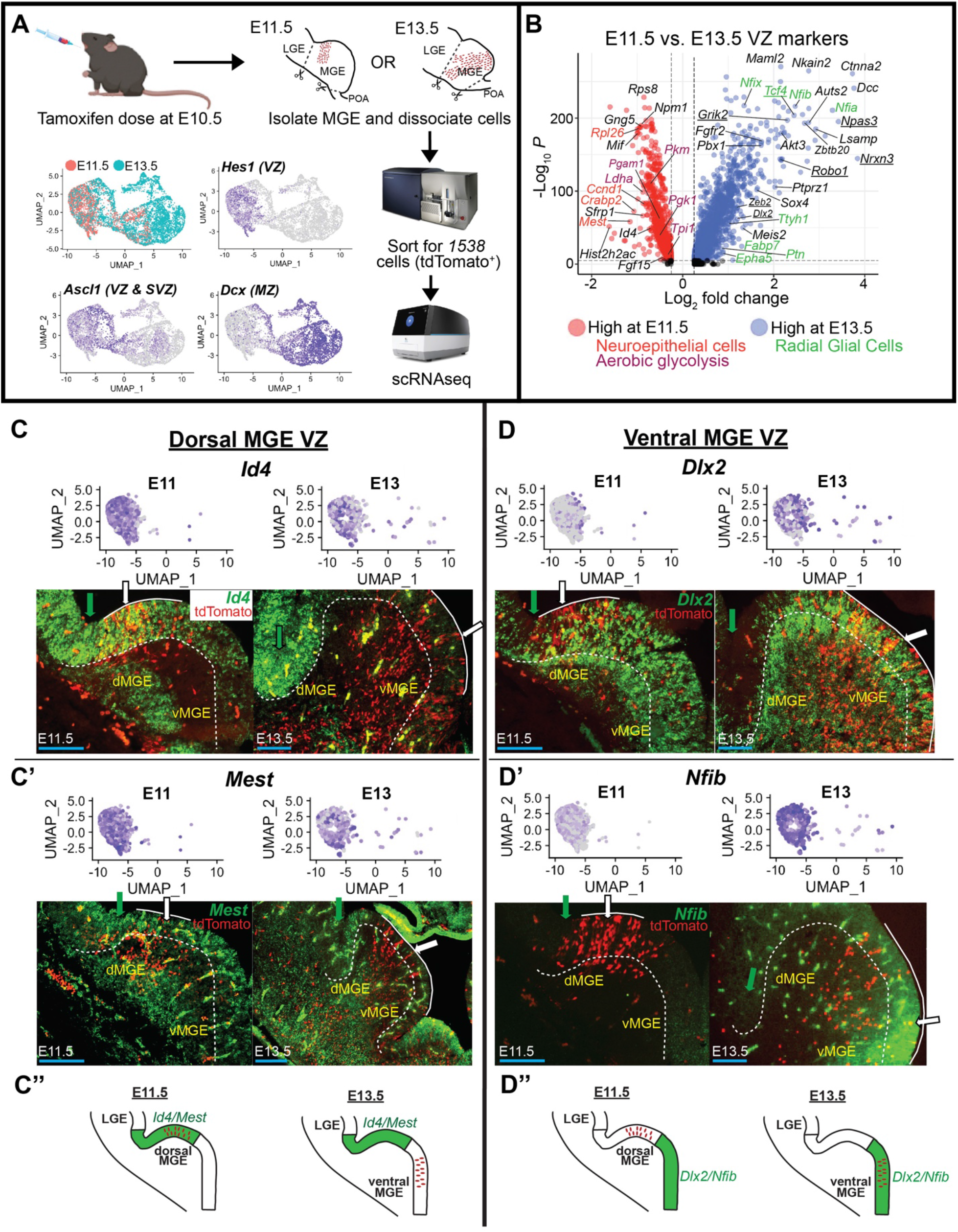
Progenitors of the 1538 lineage change gene expression and mature as they move ventrally along the MGE VZ. **(A)** Experimental paradigm of scRNAseq analysis on E11.5 and E13.5 *1538* lineages. Mice were given tamoxifen at E10.5, MGE tissue was isolated and dissociated from E11.5 or E13.5 embryos, and MGE cells were FAC sorted for the 1538 lineage using tdTomato and then processed for scRNAseq. UMAP plot of FACS purified (tdTomato^+^) E11.5 and E13.5 1538 MGE lineages; color-coded by age (top left). Feature plots showing VZ, SVZ and MZ marker expression in sorted 1538 MGE lineages (*Hes1* (top right), *Ascl1* (bottom left), and *Dcx* (bottom right)). Number of cells: 3064 (E11.5) and 4721 (E13.5); Mean expressed genes per cell: 4614.\ **(B)**Volcano plot showing top genes expressed in E11.5 (negative Log_2_ fold change, red plots) and E13.5 (positive Log_2_ fold change, blue plots) 1538 MGE VZ cells. Markers of neuroepithelial cells (red label) and aerobic glycolysis (purple label) are highly expressed in E11.5 VZ cells while radial glial cell markers are highly expressed in E13.5 VZ cells (green label). **(C, D)** Feature plots of E11.5 and E13.5 1538 VZ cells and fluorescent ISH (green) on E11.5 and E13.5 MGE co-stained with the1538 lineage marker (tdTomato^+^, red) showing the expression of dorsal MGE VZ markers, *Id4* (C) and *Mest* (C’), and ventral MGE VZ markers, *Dlx2* (D) and *Nfib* (D’). (C”, D”) Illustration of progenitors from the E10.5 *1538* lineage moving from the dorsal MGE VZ that expresses *Id4* and *Mest* at E11.5 (C”) into the ventral MGE VZ that expresses *Dlx2* and *Nfib* at E13.5 (D”). Green arrow, MGE/LGE sulcus. White arrow, approximate center of tdTomato^+^ *1538* lineage within the VZ. White solid line, distribution of *1538* lineages in the VZ. White dash line, MGE SVZ and MZ boundary. Blue scale bar, 100 uM.

We then performed differential expression (DE) analysis on E11.5 vs E13.5 VZ (*Hes1*^+^) cells. ∼1060 genes were enriched at E11.5 and ∼2220 genes were enriched at E13.5 (table S1). At E11.5, among the top 100 genes were markers for neuroepithelial cells (*Rpl26, Ccnd1, Crabp2, Mest*) or genes involved in aerobic glycolysis (*Pkm, Pgam1, Ldha, Pgk1, Tpi1*), a property of immature progenitors (*11*) (Fig 3B). At E13.5, the top 100 genes were markers of radial glial cells (*Nfia/b/x, Tyh1, Ptn, Fabp7, Epha5*) and genes involved in cortical interneuron (CIN) development (*Tcf4, Grik2, Npas3, Nrxn3, Zeb2*) (Fig 3B).

We verified the expression patterns of some of these genes using *in situ* hybridization (ISH). For instance, we used fluorescent ISH to verify the E11.5-to-E13.5 changes in VZ expression. The location of the 1538 lineage was identified by tdTomato expression (Fig. 3C, D). At E11.5, VZ expression of *Id4* and *Mest* were high and restricted to the dorsal MGE where they overlapped with tdTomato (Fig. 3C). By contrast, E13.5 *Id4* and *Mest* expression were reduced, particularly in the ventral MGE, where tdTomato cells had moved (Fig. 3C). On other hand, at E11.5, *Dlx2* and *Nfib* expression were low in the dorsal MGE (especially the MGE/LGE sulcus), but high in the VZ of the E13.5 ventral MGE where they overlapped with tdTomato expression (Fig. 3D). ISH also identified TFs, most known to regulate MGE development (*2*), that are temporally and regionally expressed in the MGE progenitors (*Lhx8, Maf, Mafb, Meis2, Nfib, Nr2f1, Otx2, Pbx1*) (fig. S6).

Thus, 1538 lineage progenitors, originating from the MGE/LGE sulcus, change gene expression as they move caudoventrally within the VZ. Moreover, based on gaining expression of *Dlx2* (a neurogenic gene enriched in the SVZ (*12*)), and the E11.5-to-E13.5 switch between neuroepithelial to radial glial markers (Fig. 3B), we postulate that VZ progenitors become more mature as they move ventrally.

### Progenitors of the 1538 lineage produce different types of neurons as they move caudoventrally along the MGE VZ

Next, we studied temporal RNA expression changes in neurons generated by the 1538 lineage by performing DE analysis on E11.5 vs E13.5 *Dcx*^+^ MZ cells (Fig. 4A). We found ∼1300 RNAs enriched in E11.5 MZ and ∼2100 RNAs enriched in E13.5 MZ cells (table S2). At E11.5, markers of projection neurons were enriched (*Meis2* and *Mest*) (Fig. 4A, B, fig. S6). On the other hand, at E13.5, markers of cortical interneuron (CINs) were enriched (*Calb1, Grik2, Maf, Mafb, Nfib, Npas3, Npy, Nrg3, Nrxn3, Sox6, Sst, Robo1*, and *Zeb2* (Fig. 4A, fig. S6)). Some projection neuron RNAs were present in the MGE SVZ and MZ (*Lhx8* and *Zic1*) or MZ only (*Gbx2*) at both ages (fig. S6).

**Fig. 4.**
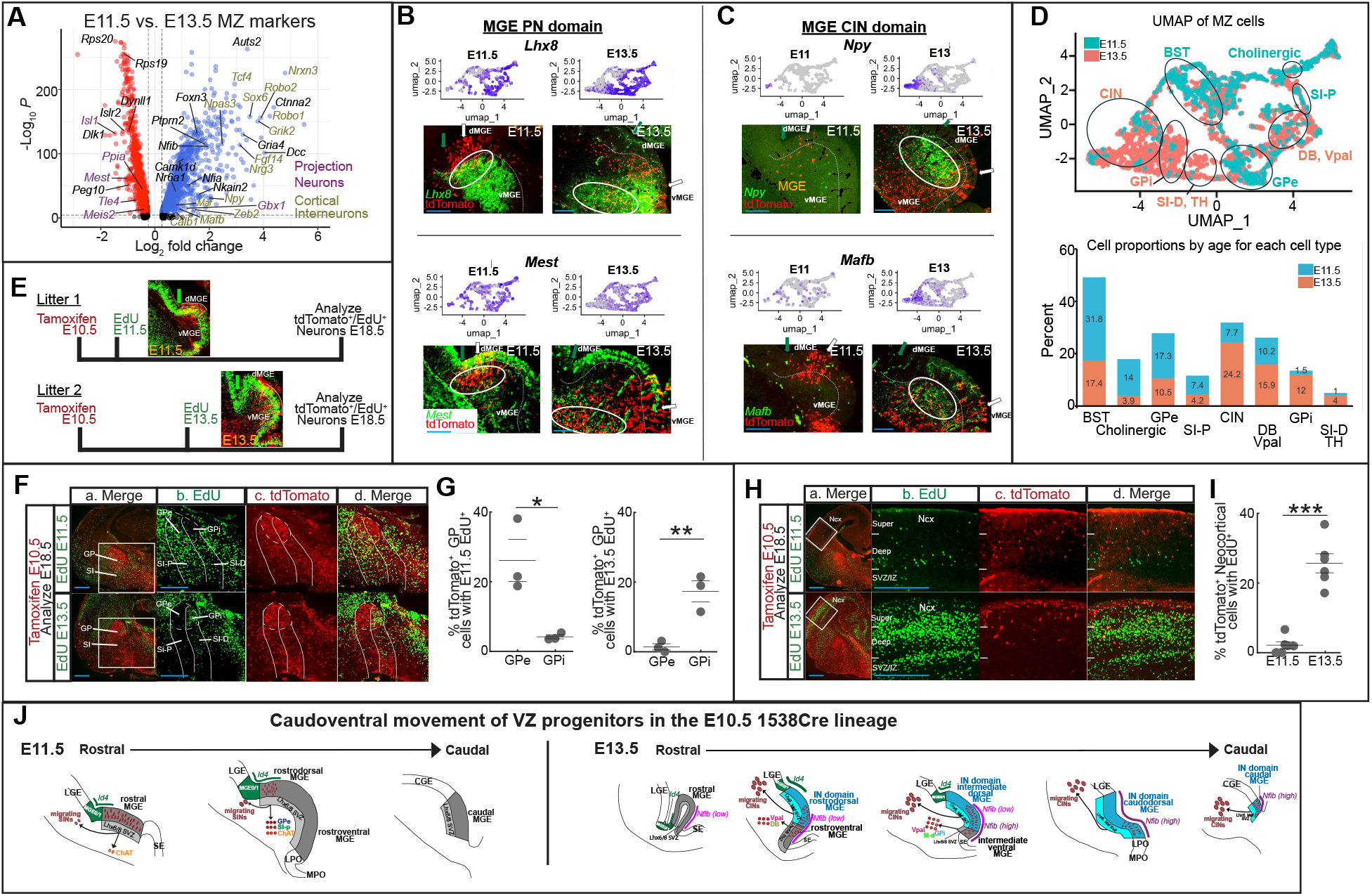
The 1538 lineage produces different neurons as they move ventrally along the MGE VZ. **(A)** Volcano plot showing top MZ cell genes expressed in E11.5 1538-lineage (negative Log_2_ fold change, red plots) and E13.5 (positive Log_2_ fold change, blue plots). Projection neurons markers (purple label) are expressed at both ages while CIN markers are primarily expressed at E13.5 (light green label). **(B, C)** Feature plots of E11.5 and E13.5 1538 MZ cells and fluorescent ISH (green) on E11.5 and E13.5 MGE co-stained with the 1538 lineage (tdTomato^+^, red) showing the expression of MGE MZ markers. *Lhx8* (C, top panels) and *Mest* (C, bottom panels) largely label immature projection neurons whereas *Npy* (D, top panels) and *Mafb* (D, bottom panels) largely label immature cortical interneurons. Green arrow, MGE/LGE sulcus. White arrow, center of 1538 VZ lineage. White dash line, MGE SVZ and MZ boundary. White ovals, tdTomato^+^ and *Lhx8*^+^, *Mest*^+^, *Npy*^+^, or *Mafb*^+^ coexpression. Blue scale bar, 100 uM. **(D)** Re-clustered UMAP plot of 1538 MZ cells (left panel) from *Dcx*^+^ cells in (Fig. 3A) (color-coded by age). Number of cells: 4047; Mean expressed genes per cell: 4629. Cell-type identity was assigned for each UMAP region based on cluster analysis in (fig. S7A) and using known expression markers (table S4). Histogram showing proportion of E11.5 and E13.5 MZ cells for each cell type (right panel). **(E)** Experimental paradigm of tamoxifen/EdU experiment. In litter 1, neurons born at E11.5 (labelled with EdU, green) from 1538 progenitors (colabelled with tdTomato^+^, red) arise from the dorsal MGE (inset: E11.5 MGE). In litter 2, neurons born at E13.5 (labelled with EdU, green) from 1538 progenitors (colabelled with tdTomato^+^, red) arise from the ventral MGE (inset: E13.5 MGE). White solid line, distribution of 1538 lineages in the VZ. **(F)** Merged images (Column a) of EdU (green) from injected at E11.5 (top row) or E13.5 (bottom row) and 1538 lineage (red, tdTomato^+^) in E18.5 coronal sections. White box in Column a indicates pallidal regions magnified in columns b-d. Magnified views: Column b: EdU; Column c: tdTomato; Column d: merged EdU and tdTomato. GPe, globus pallidus externa; GPi, globus pallidus interna; SI-d, substantia innominata diagonal; SI-p, substantia innominata pallidal. Blue scale bar, 200 uM. **(G)** Comparing birthdate of GPe and GPi in 1538 lineage. Percentage of tdTomato^+^ cells colabeled with EdU injected at E11.5 (left panel) or E13.5 (right panel) in E18.5 GPe and GPi (n=3 for each birthdate). *, p<0.05; **, p<0.01. **(H)** Birthdating of CINs in 1538-lineage. Merge images (Column a) of EdU (green) from injected at E11.5 (top row) or E13.5 (bottom row) and 1538 lineage (red, tdTomato^+^) in E18.5 coronal sections. White box in Column a indicates neocortical area magnified in columns b-d. Magnified views: Column b: EdU; Column c: tdTomato; Column d: merged EdU and tdTomato. Deep, layers V and VI; Ncx, neocortex, Super, layers I thru IV; SVZ/IZ, subventricular zone/intermediate zone. Blue scale bar, 200 uM. **(I)** Percentage of tdTomato^+^ cells in E18.5 neocortex co-labeled with EdU injected at E11.5 or E13.5 (n=6 for each birthdate). ***, p<0.001. **(J)** Schema showing that 1538 lineages (initiated with E10.5 tamoxifen; solid red circles) produce different neurons (red circles with different-colored cores) as they move in a caudoventrally along the MGE VZ. The proposed growth zone is green and expresses *Id4*. Left panel: At E11.5, 1538 progenitors occupy rostrodorsal MGE positions that express *Lhx6* and *Lhx8* in the SVZ (grey); we propose that these produce SIN (striatal interneurons), ChAT^+^ (cholinergic), GPe (globus pallidus externa), and SI-p (substantia innominata pallidal) neurons. Right panel: At E13.5, 1538 progenitors move caudoventrally away from the *Id4*^*+*^ growth zone (green label) and towards the *Nfib*^*+*^ zone (purple label). At E13.5, the dorsal *Lhx6*^+^ but *Lhx8*^-^ region (blue) promotes interneuron (CIN) production. The 1538 lineage continues to produce different types of projection neurons in the ventral MGE: DB (diagonal band), GPi (globus pallidus interna), SI-d, substantia innominate diagonal, and Vpal (ventral pallidum) neurons.

We then reclustered E11.5 and E13.5 *Dcx*^+^; tdTomato^+^ 1538 lineage MZ cells using a gene list of 52 known regulators and markers of projection neuron and cortical interneuron development (table S3) and mapped them on the same UMAP plane to define different types of neurons present at these ages (Fig. 4D, figs. S6, S7). Clusters were identified based on their combinatorial expression of RNAs that mark specific cell types (table S4). The proportion of E11.5 and E13.5 cells were calculated for each cluster/cell type. For example, E11.5 neurons included bed nucleus stria terminalis (BST), cholinergic, and external globus pallidus (GPe) (Fig. 4D). In contrast, E13.5 neurons included CINs, internal globus pallidus (GPi), and ventral pallidum (Vpal) neurons (Fig. 4D).

When comparing *Lhx6* and *Lhx8* expression, we noted that at E11.5, they shared indistinguishable domains in the MGE SVZ and MZ (Fig. 4J, fig. S6, S8) (*13*). However, by E13.5, *Lhx8* is repressed throughout most of the dorsal MGE SVZ (Fig. 4J, fig. S6, S8); its expression remains in the MGE MZ. Furthermore, the *Lhx8*^-^ zone expresses CIN markers (*Calb1, Maf, Mafb, Npy, and Sst*) (Fig 4C and J, figs. S6, S8). Thus, there is a major molecular reorganization of the MGE between E11.5 and E13.5. Given that, we examined the position of 1538 lineage cells with respect to *Lhx8, Mafb, Mest*, and *Npy* (using two-color fluorescent ISH; Fig 4B, C).

At E11.5, the 1538 lineage SVZ and MZ cells expressed RNA markers of projection neurons but not CINs. For instance, *Lhx8* and *Mest* were expressed in 1538 lineage dorsal MGE SVZ and MZ cells (Fig. 4B). By contrast, there was no *Npy* and *Mafb* expression in this lineage (Fig. 4C). This finding leads to the notion that at E11.5, when the 1538 lineage is in the rostrodorsal MGE, it primarily produces projection neurons. Expression of projection neuron markers, *Pbx1* and *Zic1*, support this hypothesis (fig. S6). However, by E13.5, CIN markers (*Mafb* and *Npy*) are expressed in the 1538 lineage in the *Lhx8*^-^ domain (Fig. 4C). We propose that this region largely generates CINs.

Our scRNAseq data implies that different types of neurons are being produced by the 1538 lineage over time during its movement within the MGE VZ. To test this hypothesis, we performed an EdU pulse chase. Two sets of mothers were given tamoxifen at E10.5 (Fig. 4E). One set was then injected with EdU at E11.5 while the other set was injected with EdU at E13.5. Embryos were analyzed at E18.5. tdTomato^+^, EdU^+^ cells from E11.5 EdU injected embryos marked neurons born at E11.5 while the 1538 lineage was in the dorsal MGE. On the other hand, tdTomato^+^, EdU^+^ cells from E13.5 injected embryos marked neurons born at E13.5 when the 1538 lineage was in the ventral MGE.

The 1538 neurons born around E11.5 occupied dorsal MGE MZ regions such as the GPe and the pallidal substantia innominata (SI-P) but generated very few CINs (Fig. 4F-I). Conversely, 1538 neurons born at E13.5 occupied ventral MGE MZ regions such as the GPi and the diagonal substantia innominata (SI-D) (Fig. 4F, G). CINs are robustly generated at this age (Fig. 4H, I). Thus, we hypothesize that progenitors of the E10.5 1538 lineage first produce GPe and SI-P neurons from the dorsal MGE and then change gene expression as they move ventrally along the MGE VZ to promote the production of GPi and SI-D neurons and CINs (Fig. 4J).

### Nfia/b dcKO regulate CIN numbers and subtype specification

*Nfia* and *Nfib* expression increased at E13.5, a stage when the progeny of *1538* progenitors moved into the ventral MGE (Fig 3B, D; figs. S6, S9). To study their function, we made conditional *Nfia* and *Nfib* double mutants (*Nfia/b* dcKO) by crossing them with *BAC-Nkx2*.*1-Cre* mice (*14*) and analyzed them at E15.5, P1, and P30 (Fig. 5, fig. S9). At P30, the mutants had ∼20% increase in tdTomato^+^ cell density in the neocortex (Fig. 5A). *Sst*^+^ CIN density increased by ∼25%, whereas PV^+^ CIN (Parvalbumin) density decreased by ∼25% (Fig 5B, C). Surprisingly, *Vip*^+^ CIN (Vasoactive Intestinal Peptide) density in MGE-derived CINs was increased (Fig 5D, fig. S9). This was unexpected because *Vip*^*+*^ CINs are CGE derived (*15*). We next determined if the increase in tdTomato^+^, *Sst*^+^ and *Vip*^+^ numbers occurred earlier than P30. TdTomato^+^, *Sst*^+^, and *Vip*^+^ cell densities were increased in the neocortex at P1 (Fig. 5E-G). At E15.5, the tdTomato^+^ and Sst^+^ increase occurred in the ventral and caudal neocortex (fig. S9). Moreover, we did not observe a change in tdTomato^+^ densities in the GP and Vpal. In summary, *Nfia* and *Nfib*, whose expression increases at E13.5 when it is largely restricted to the VZ, regulate CIN numbers and subtype identity, but does not regulate projection neuron production. Note that E13.5 is a peak age when CINs are produced (Fig. 4H, I) and when 1538 progenitors arrive in the ventral MGE (Fig. 1) where *Nfia* and *Nfib* expression are highest (Fig. 5H).

**Fig. 5.**
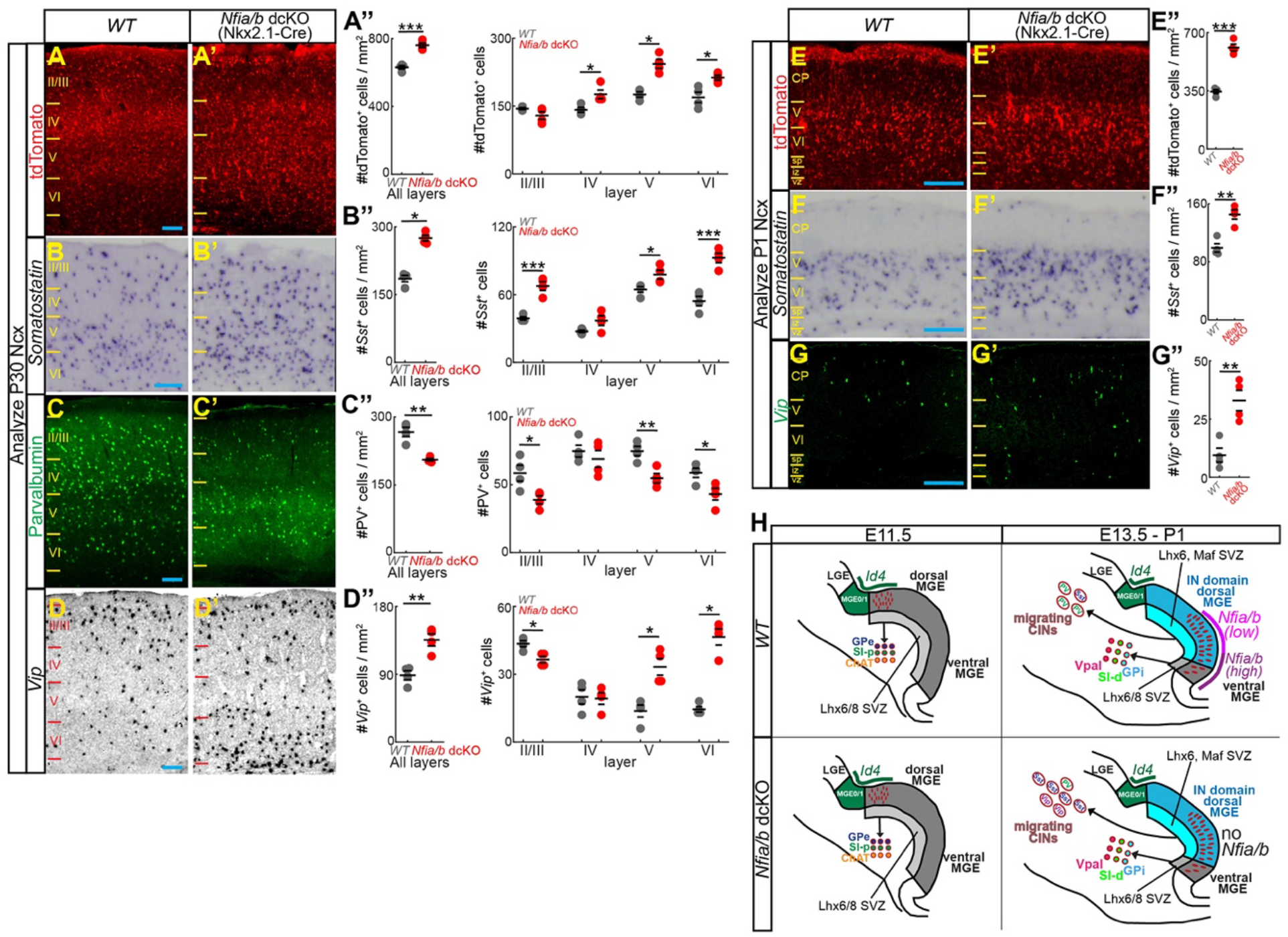
*Nfia* and *Nfib* temporally regulate the number and identity of CINs. **(A-D)** tdTomato immunohistochemistry (IHC) (A, A’), *Somatostatin* ISH (B, B’), Parvalbumin IHC (C, C’), and *Vip* ISH (D, D’) on P30 somatosensory neocortex from wildtype (A-D) and *Nfia/b* dcKO (A’-D’) mice. tdTomato^+^ (A”), *Somatostatin*^+^ (B”), Parvalbumin^+^ (C”), and *Vip*^+^ (D”) densities in all layers (left panels) or in individual neocortical layers (right panels) (n=4 for each genotype). **(E-G)** tdTomato IHC (E, E’), *Somatostatin* ISH (F, F’), and *Vip* ISH (G, G’) on P1 somatosensory neocortex from wildtype (E-G) and *Nfia/b* dcKO (E’-G’) mice. tdTomato^+^ (E”), *Somatostatin*^+^ (F”), and *Vip*^+^ (G”) densities in all neocortical layers (n=4 for each genotype). CP, cortical plate; IZ, intermediate zone; sp, subplate; VZ, ventricular zone. Error bars, SEM. *, p<0.05; **, p<0.01, ***, p<0.001. Blue scale bar, 100 uM. **(H)** Repattering of the MGE from E11.5 to E13.5; creation of an interneuron generating domain. At E11.5, the MGE has little *Nfia* and *Nfib* expression but has broad *Lhx6* and *Lhx8* SVZ expression (grey) to produce projection neurons. However, by E13.5, the region of *Lhx8* expression is largely replaced by strong *Nfia* and *Nfib* expression (pink/purple) in the dorsal MGE to create an interneuron generating domain (blue, *Lhx6*^+^/*Maf*^+^). At E13.5, CIN production and subtype identities are affected in *Nfia/b* dcKO as illustrated. Defects in projection neurons have not been detected in the mutant. Movement of the *1538* lineage from E11.5-E13.5 is shown (red dots in VZ). The putative growth zone at MGE/LGE sulcus (MGE0/1) is shown in green as is the expression of *Id4*. Interneurons: CIN, cortical interneurons; PV, parvalbumin; Sst, somatostatin; Vip, vasoactive intestinal peptide. Projection neurons: GPe, globus pallidus externa; GPi, globus pallidus interna; SI-d, substantia innominata diagonal; SI-p, substantia innominata pallidal; Vpal, ventral pallidum.

## Discussion

The current dogma of CNS regional molecular patterning of the VZ is that it has sharp boundaries of TF expression that specify fixed regional subdomains (*1-3*). Here we challenge this conception for early stages of mouse MGE development (E9.5-E12.5). Note that the current MGE model was based on TF expression at E13.5 (*1*,*2*). Here we show that MGE VZ cells generated in a rostrodorsal growth zone (Septal/MGE (see fig. S1) and LGE/MGE sulcus) progressively moved caudoventrally. We propose that they are displaced by newly generated progenitors until ∼E13 (Figs. 1 and 2; figs. S2 and S4). This phenomenon may share similarities with other developmental growth zones (e.g. limb progress zone, somitogenesis, Hensen’s node) (*16-18*). We hypothesize that the embryonic brain may have other regions analogous to the MGE growth zone. For instance, fate-mapping from the *Fgf8/17*^+^ rostral telencephalic/septal patterning center generated clones that dispersed tangentially (planarly) in the cortical VZ (*19*). In all, we propose that MGE morphogenetic patterning connects its growth to regional patterning and cell type specification.

MGE TF boundaries are maintained during this process (Fig. 3); thus, the moving progenitors must change their transcriptional profile as they move into adjacent domains, implying that they dynamically degrade some TFs (e.g. *Id4, Lhx8*), while inducing the expression of other TFs (e.g. *Nfi* family). We hypothesize that progenitors may have a cell autonomous mechanism (perhaps based on timing) to accomplish this complex process, although we do not rule out that positional information (such as derived from SHH concentration) also contributes to their changes in gene expression as they move. However, cKO of *Ptch1*, a receptor for SHH, did not affect MGE VZ displacement (*1538CreERT2*, tamoxifen E10.5; data not shown), suggesting that SHH-signaling is not critical.

The mechanism of progenitor movement is likely to be caused by displacement of newly arriving progenitors. We do not think that the tdTomato labeled VZ cells actively migrate planarly in the traditional sense. The process ends around E13.5. At that stage the *1538CreERT2* lineage no longer moves away from the growth zone and their descendants primarily radially migrate toward the pial surface (Fig. 1B, C).

Over time, as the 1538 VZ lineage’s changes position, it generates different neuronal subtypes. For instance, GPe, and SI-P neurons are born at E11.5 when the 1538 lineage is in the rostrodorsal MGE, while GPi and SI-D neurons are born at E13.5 when the 1538 lineage is in the caudoventral MGE (Fig. 4F, G). This is consistent with fate-mapping and histochemical studies showing that GP regional subdivisions have distinct origins and molecular properties (table S4) (*5*,*20*). Other early generated (E11.5) cell types include parts of the BSTN, which is unexpected because their periventricular (“deep”) location would normally imply that they are produced late, and subsets of cholinergic neurons (Fig. 4D; Fig S7). Cholinergic neurons are largely generated by *Fgf8*^*+*^*/17*^+^ progenitors (*19*), which is consistent with the proximity of the rostral-most *1538*^+^ MGE growth zone to the septum (fig. S2, S4).

Timing also influences the relative generation of PNs versus CINs. At E11.5, the *1538* lineage primarily produces pallidal projection neurons, as evident by their location in the basal ganglia (Fig. 4B; fig. S6), and the expression of projection neuron markers/regulators such as *Lhx8, Meis2*, and *Mest* (Fig. 4A) (*21*). By contrast, at E13.5, while some projection neurons are still being produced, relatively more CINs are generated, based on their location (Fig. 4C; fig. S6), EdU pulse labeling, and the induction of genes (e.g *Maf, Nfia/b, Npy*) that mark/specify CINs (Fig. 3 and 4). We postulate that the induction of CIN generation is associated with repression of *Lhx8* in the MGE VZ/SVZ (compare E11.5 with E13.5; Fig. 4B). The emergence of the interneuron producing domain (Fig. 3B, 4A, C, J) coincides with the *Dlx*-mediated up-regulation of *Zeb2* which represses *Nkx2*.*1* and *Lhx8* (*22*). Temporal regulation of MGE cell fate has also been shown for *Nr2f2*, which promotes GP fate at E11.5 and *Sst*^+^ CINs at E13.5 (*4*). Overall, we hypothesize that E11.5 expression of genes such as *Lhx8, Mest*, and *Id4* promote projection neuron fate (Fig. 3C, 4B, J, fig. S6); by E13.5 these genes are downregulated in a specific part of the MGE where genes involved in CIN specification are upregulated (e.g *Dlx2, Maf, Nfia/b, Npy*)(Fig. 3D, 4C, J, fig. S6).

*Nfia* and *Nfib* temporally and autonomously regulate a switch in MGE progenitors after E11.5 by promoting PV^+^ and repressing SST^+^ CIN fate (Fig. 5). *Nfia* and *Nfib* are upregulated in the VZ at E13.5 when the *1538* VZ lineage moves into the caudoventral MGE. *Nfia* and *Nfib* also control the temporal generation of specific neuronal subtypes in the neocortex and retina (*23, 24*). While the MGE CIN phenotype in the *Nfia/b* dcKO is similar to the one in the *Mafb;Maf* cKO (*25*), we did not detect a change in *Maf* expression (fig. S9). *Nfia* and *Nfib* may regulate MGE CIN specification by repressing *Nr2f2*, since this TF promotes Sst and represses PV CIN fate (*4*).

A previous study used tCROP-seq to mutate *Nfib* and *Nfix* in the entire E12.5 ganglionic eminences and found reduced GABAergic neurons in the E15.5 striatum (*26*), whereas we found in the *Nfia/b* dcKO increased neuronal output from the MGE. Perhaps, differences in the experimental design account for these divergent results.

Surprisingly, *Nfia* and *Nfib* repress VIP^+^ CIN fate in MGE progenitors (Fig. 5). The MGE genes, *Nkx2*.*1* and *Lhx6*, also repress CGE CIN fates (*27, 28*). Note, however that *Lhx6* mutants do not ectopically produce VIP^+^ CIN (*28*). Thus, *Nfia* and *Nfib* may regulate VIP^+^ CIN fate through a pathway that is *Lhx6* independent.

In sum our novel proposal that MGE morphogenetic patterning connects its growth to regional patterning and cell type specification raises important questions as to how molecular specification of progenitors are regulated as these cells move caudoventraly along the VZ of the MGE.

## Acknowledgements

University of California, San Francisco (UCSF) cores supported this work. Sequencing was carried out at the UCSF Center for Advanced Technology, and cell sorting was carried out at the UCSF Helen Diller Family Comprehensive Cancer Center Laboratory for Cell Analysis (P30CA082103). We thank Mylinh Bernardi and Horng-ru Lin of the Gladstone Genomics Core for their assistance with the 10x Genomics scRNAseq library preparation.

## Author Contributions

Conceptualization: JSH, JLR

Methodology: JSH, KC, ASN, JLR

Validation: JSH, KC, LP

Formal Analysis: JSH, KC

Investigation: JSH, KC, LP, ASN, JLR

Resources: LJR, ASN, JLR

Writing-original draft: JSH, KC, JLR

Writing-review & editing: JSH, KC, JWCL, LJR, LP, ASN, JLR

Supervision: LP, ASN, JLR

Funding acquisition: LJR, LP, ASN, JLR

## Data and materials availability

Sequencing data will be deposited at the National Center for Biotechnology Information BioProjects Gene Expression Omnibus and will be accessible through a GEO accession number. *Nfia* flox and *Nfib* flox mice are available from Linda J. Richards (Washington University in St. Louis School of Medicine), and *1538-CreERT* mice are available from John L. Rubenstein (University of California, San Francisco). All data needed to evaluate the conclusions in the paper are present in the main text or the supplemental materials.

## Supplementary Materials

Materials and Methods

Figs. S1 to S9

Tables S1 to S4

