## Supplemental Material & Methods, and Figures for "Morphogenetic Patterning During Regional and Cell Type Specification in the Embryonic Basal Ganglia"

#### Materials and Methods

Animals and Brain Isolation

We used the following mouse strains: *tdTomato*$^{lox/+}$ *(Ai14)* Cre-reporter (*6*), *Nfia*$^{lox/+}$ (24), *Nfib*$^{lox/+}$ (*29*), *Nkx2.1-Cre* (*14*), and *1538-CreERT* (*5*). All strains were on a mixed C57BL/6 and CD1 background. Animals were housed in a vivarium with a 12 h light/12 h dark cycle. Postnatal animals used for experiments were kept with their littermates. Males and females were used in all experiments. All animal care and procedures were performed according to the University of California at San Francisco Laboratory Animal Research Center guidelines. Mice were anesthetized with intraperitoneal avertin (0.015 ml/g of a 2.5% solution) followed by opening of the chest cavity and perfused transcardially first with PBS and then with 4% PFA. The brain was then removed, followed by overnight fixation in 4% PFA, 2 days of cryoprotection with 30% sucrose, and microtome sectioning (coronal, 40 µm).

For timed pregnancies, noon on the day of the vaginal plug was counted as embryonic day (E) 0.5. For EdU experiments, pregnant mice were injected intraperitoneally with a single dose of 50 mg/kg body weight (from a 10 mg/ml stock), and brains were harvested at E18.5 (Fig. 4, fig. S7). For tamoxifen experiments, pregnant mice were administered 1 mg/40 kg body weight (from a 10 mg/ml stock dissolved in corn oil) by oral gavaging with silicon-protected needles. Mice were euthanized with $CO_2$ inhalation followed by cervical dislocation and the brains from the embryos were isolated. Brains used for *in situ* hybridization and immunohistochemistry were fixed overnight in 4% paraformaldehyde (PFA), transferred to 30% sucrose overnight for cryoprotection, and then OCT embedded for cryosectioning. Sections were cut coronally at 20 µm.

Immunohistochemistry (IHC)

IHC on embryonic brains was performed on slide-mounted cryosections. These were rinsed in phosphate-buffered saline (PBS), permeabilized in PBST (PBS with 0.1% Triton X-100), blocked in 5% normal serum/PBST, incubated in primary antibody overnight or for two days at 4°C, washed in PBST, incubated with secondary antibody for 1-2 h (at room temperature) and washed in PBS. For IHC on postnatal brains, staining was carried out on free-floating sections as described previously (*30*). EdU staining was performed after the secondary antibody incubation and labeled with a Click-iT EdU Alexa Fluor Imaging kit (Catalog #C10337, Molecular Probes). We performed all IHC on $n \geq 3$ biological replicates for each control and mutant.

Antibodies

Antibodies used were rabbit DsRed (1:500, Catalog #632496, Clontech), rat RFP (1:500, Catalog #5F8, ChromoTek), rat Lamp5, clone 34.2 (1:100, Catalog #14-9778-80, ThermoFisher), rabbit nNOS (1:200, Catalog #61-7000, Life Technologies), rabbit NPY (1:750, Catalog #22940, Immunostar), rabbit pH3 (1:500, Catalog #06-750, Upstate), mouse Parvalbumin (1:300, Catalog #MAB1572, Swant Swiss Abs), mouse Reelin (1:300, Catalog #MAB5364, Millipore), rabbit Tyrosine Hydroxylase (1:500, Catalog #AB152, Millipore), and Alexa-conjugated secondary antibodies (1:300, Molecular Probes). Sections were cover slipped with Vectashield (Vector labs).

*In Situ* Hybridization (ISH)

We performed ISH on a minimum of n=3 biological replicates for each control and mutant. In each case, a rostrocaudal series of at least ten sections was examined. ISH were performed using digoxigenin-labeled riboprobes as described previously (*19*). Digoxigenin-labeled riboprobes were made against *Maf*, *GFP*, *Somatostatin*, and *Vip*.

Dual RNAscope (Fluorescent *In Situ* Hybridization) and IHC Labeling

Fluorescent ISH was performed according to the manufacturer's protocol for the multiplex fluorescent kit (ACD, version 2, Cat. 323100) with the following modifications: 1) sections were treated with 1x Target Retrieval Reagent at ~60$^{o}$C for 5 minutes and 2) were later treated with Protease IV for 30 minutes for P30 sections or with Protease Plus for 10 minutes for all other sections at 40$^{o}$C. The following RNAscope probes were used: Mm-Dlx2 (Cat. 555951), Mm-Id4 (Cat. 447861), Mm-Lhx6 (Cat. 422791), Mm-Lhx8 (Cat. 515101), Mm-Mafb (Cat. 438531), Mm-Mest (Cat. 405961), Mm-Mfge8 (Cat. 408771), Mm-Nfib (Cat. 586511), Mm-Npy (Cat. 313321), Mm-Ntrk1 (Cat. 435791), Mm-Ptch1 (Cat. 402811), and Mm-Vip (Cat. 415961). After ISH, IHC with the rabbit DsRed antibody (see above) and EdU labeling were performed. Lastly, slides were mounted with Vectashield mounting media containing DAPI (Cat. H-1800).

Slice Culture

Pregnant mice were euthanized with $CO_2$ inhalation followed by cervical dislocation. Embryonic brains were dissected in cold Hanks buffered saline solution (HBSS) containing 2.5 mM Hepes, 30 mM glucose, and 4mM $NaHCO_3$, and kept on ice until being embedded for coronal sectioning in 4% low melt agarose. Brains were sectioned at a 250 μm thickness with a Leica VT1200S microtome (Leica Microsystems). Sections were then transferred to 8.0 nucleopore track-etch membranes (Cat. 110414, Whatman) floating on DMEM/F12 medium containing N2, B27, 10% FBS, 10 ng/ml Fgf2, and 20 ng/ml Egf. Sections were allowed to recover by incubating it in 95% $O_2$ and 5% $CO_2$ at 37°C for 1 hour. Time in slice culture begins after this recovery period. For measuring the amount of movement of tdTomato$^{+}$ cells in slice cultures (fig. S3), dorsal-ventral position of tdTomato$^{+}$ cells in the VZ/SVZ were measured relative to the length from the MGE/LGE sulcus to the MGE/POA sulcus. These positions were plotted, and the normalized median positions were calculated for each time point.

Single Cell RNA-seq Sample and Library Preparation

For single-cell collection and dissociation, MGE tissues were dissected and pooled from E11.5 or E13.5 embryos (N= 3 or 4 embryos from 2 litters) in Earle's Balanced Salt Solution (EBSS) at room temperature, and then incubated with a prewarmed solution of Papain (Worthington; prepared according to manufacturer's protocol) for 10 minutes at 37°C. After Papain incubation, tissues were washed multiple times with ice cold EBSS (Thermo Fisher), gently triturated, and then treated with an albumin ovomucoid inhibitor mixture (Worthington; prepared according to manufacturer's protocol). Dissociated MGE cells were resuspended in ice-cold EBSS containing 1% BSA and filtered (30 μm MACS SmartStrainer) into new conical tubes. MGE cells were loaded into a FACS Aria II sorter (UCSF, Laboratory for Cell Analysis core) to isolate 1538$^{+}$ lineages by gating for tdTomato$^{+}$, DAPI$^{-}$ cells. Cell concentration and viability were confirmed using a Countess II automatic hemocytometer before loading onto a 10X Genomics single-cell preparation platform, following the manufacturer's instructions to target 10,000 cells for recovery per sample. Library preparation was performed by the Gladstone Institutes Genomics Core, using 10X Chromium Single 3' Reagent Kits Version 3.1 (10X Genomics). The library length and concentration were quantified using a Bioanalyzer (Agilent) according to the manufacturer's protocol. Sequencing was performed on the Illumina NovaSeq (UCSF, Center for Advanced Technology core).

Cell Ranger Processing

Raw single-cell RNA-sequencing data were processed on a Linux server using **Cell Ranger v7.0.1** (10x Genomics). FASTQ files from three datasets (1538TE10H11, 1538TE10H13, and JH4327_01_TE10H13) were processed independently with the *cellranger count* pipeline. Reads were aligned to the mouse reference genome (mm10) using the 10x Genomics reference **refdata-gex-mm10-2020-A**. Alignment, barcode processing, UMI collapsing, and gene expression quantification were performed using default Cell Ranger parameters. The resulting filtered gene-by-cell count matrices, barcode annotations, and quality control metrics were inspected and used as input for all downstream analyses in R. Filtered gene-by-cell count matrices generated by Cell Ranger were imported into R (v4.3.3) and analyzed using **Seurat** (v5.0.3). The integrated dataset comprised three single-cell RNA-seq libraries: **1538TE10H11 (n= 3064 cells, post QC), 1538TE10H13 (n= 1374 cells, post QC)**, and **JH4327_01_TE10H13 (n= 3347 cells, post QC).**

Quality Control

Removal of non-MGE contaminants. Preliminary clustering and marker gene inspection identified contaminating cell populations corresponding to the preoptic area (POA) and septum—regions adjacent to the medial ganglionic eminence (MGE) that are frequently captured during embryonic dissections—in the 1538TE10H11 and 1538TE10H13 datasets. These populations were removed using a two-stage computational strategy combining marker-based preselection with supervised classification. Putative POA cells were identified based on high expression of canonical POA markers (*Igfbp5*, *Dbx1*, and *Hmx3*), defined as cells within the upper quantile of the non-zero expression distribution for at least one marker. Septal cells were identified analogously using enrichment of Spry1, Fgf8, and Zic4. Differential expression analysis confirmed transcriptional profiles consistent with POA and septal identities. To refine classification, support vector machine (SVM) models were trained using the scAnnotatR framework, with marker-defined cells used as training labels. Both POA and septum classifiers achieved high discriminative performance (area under the ROC curve > 0.95). Cells classified as POA, septum, or ambiguous (POA/septum) were excluded from downstream analyses, yielding a curated MGE reference population. The filtered Seurat object was saved and used for all subsequent analyses. The JH4327_01_TE10H13 dataset showed no detectable POA or septal contamination and was integrated directly with the curated datasets. Identical quality-control thresholds and analysis parameters were applied uniformly across all datasets. Cells with fewer than 1,000 UMIs or more than 20% mitochondrial reads were excluded.

Dataset integration, Dimensionality Reduction, and Clustering

All datasets were normalized using SCTransform, which models sequencing depth–dependent technical variation using a regularized negative binomial framework implemented via glmGamPoi. Data integration was performed using Seurat's SCT-based workflow. Briefly, 3,000 integration features were selected across datasets using SelectIntegrationFeatures, followed by preparation for SCT integration with PrepSCTIntegration. Integration anchors were identified using FindIntegrationAnchors (normalization.method = "SCT"), and datasets were integrated using IntegrateData with the default anchor weighting (k.weight = 100) to account for unequal dataset sizes. The integrated expression matrix was subjected to principal component analysis (PCA) using RunPCA. The number of retained components was determined empirically based on variance explained and inspection of scree plots, and 25 principal components (PCs) were selected for downstream analyses. Nonlinear dimensionality reduction was performed using Uniform Manifold Approximation and Projection (RunUMAP) with the selected PCs. A shared nearest-neighbor graph was constructed in the integrated space using FindNeighbors, and unsupervised clustering was performed using the Louvain algorithm via FindClusters on the integrated SNN graph. Integration quality was assessed by visualization of UMAP embeddings colored by dataset and developmental stage, as well as by inspection of known marker gene expression using feature and violin plots, confirming effective alignment of shared cell states while preserving biologically meaningful differences across developmental stages. The developmental organization and orientation of the UMAP embedding were interpreted using expression gradients of *Hes1*, *Ascl1*, and *Dcx*, which mark ventricular zone progenitors (VZ), subventricular zone neurogenic intermediates (SVZ), and postmitotic mantle zone neurons (MZ), respectively.

Reclustering of the MZ cells

To achieve higher-resolution characterization of MZ neuronal diversity, MZ cells were reintegrated and reclustered using a gene-set–restricted strategy focused on projection neuron and cortical interneuron identity. A curated set of 52 projection neuron and cortical interneuron associated genes was used to subset the expression matrix prior to reintegration. These genes were selected based on known roles in projection neuron and cortical interneuron specification, subtype identity, and maturation and is included in table S3. MZ cells were first subset from the globally integrated Seurat object based on cluster identity and then split by dataset. Each dataset was independently normalized using SCTransform, after which reintegration was performed using the SCT-based integration workflow (SelectIntegrationFeatures, PrepSCTIntegration, FindIntegrationAnchors, and IntegrateData; 3,000 integration features). Importantly, reintegration was carried out only on the restricted projection neuron and cortical interneuron gene set, thereby emphasizing transcriptional variation relevant to neuronal subtype identity while minimizing contributions from unrelated gene programs. Dimensionality reduction was performed on the reintegrated data using principal component analysis (RunPCA; 25 PCs), followed by UMAP embedding with parameters tuned to enhance local structure and cluster separation (RunUMAP, dims = 1–25, min.dist = 0.01). Graph-based clustering was then performed using FindNeighbors and FindClusters (resolution = 0.8), yielding a refined MZ clustering with increased separation of projection neuron and cortical interneuron subtypes compared to the original global UMAP. After reclustering, the resulting PCA and UMAP embeddings, as well as updated cluster identities, were transferred back to the corresponding MZ subset of the original Seurat object. This step preserved the full expression matrix across all genes while enabling visualization and downstream analyses (e.g., heatmaps, FeaturePlots, and differential expression) in the context of the refined MZ embedding. Reclustering outcomes were compared to the original clustering using alluvial analyses, demonstrating strong correspondence with the global structure while revealing additional resolution within the neuronal compartment.

Image Acquisition and Analysis

Fluorescent IHC images were taken using a Coolsnap camera (Photometrics) mounted on a Nikon Eclipse 80i microscope using NIS Elements acquisition software (Nikon). Brightfield ISH images were taken using a DP70 camera (Olympus) mounted on an Olympus SZX7 microscope. Brightness and contrast were adjusted, and images merged using ImageJ software.

Cell Counting

For $pH3^+$ density measurements (Fig. 2A, fig. S5), the dorsal-ventral position of $pH3^+$ cells in the VZ were measured relative to the distance between the MGE/LGE sulcus to the MGE/POA sulcus. These positions were then binned into 20 equally spaced bins. Density was calculated by tallying the total number of $pH3^+$ cells for each bin.

For assessing EdU, Lamp5, nNos, Npy, Parvalbumin, Reelin, *Sst*, tdTomato, and *Vip* cell densities (Fig. 4, 5; fig. S8), 10× images were taken from the somatosensory cortex (for postnatal ages) or from the neocortex (for embryonic ages); pictures were taken of two or three nonadjacent sections and from both hemispheres for each replicate. For quantification, a box of a defined area was drawn over the region where we counted cell numbers. Cell density was calculated by the number of cells expressing a marker divided by the box area. For lamination counts, we used DAPI as a guide for subdividing the neocortex into its layers.

To count EdU and tdTomato densities in the globus pallidus (GP), an oval was drawn around the GP. GP cells were anatomically defined in the mantle zone of coronal MGE sections (fig. S2). The GP was subdivided into GPe and GPi based on the relative positions (*20*). Percent $tdTomato^+$ cells containing EdU labelling within each GP portion was calculated by dividing the number of cells labelled both with EdU and tdTomato by the total number of cells expressing tdTomato.

Quantification and Statistical Analysis

All bar graphs show mean ± SEM. All statistical analyses were performed using GraphPad Prism (version 7), and a p-value of $< 0.05$ was considered significant. The specific numbers of replicates (n) for each experiment can be found in the figure legends.

For all cell counts involving IHC and ISH, we used the cell counter plug-in in FIJI software. All IHC and ISH statistical analyses were carried out on SPSS15 software and Microsoft Excel. All data points were found to lie within a normal distribution using a Shapiro-Wilk test and were therefore suitable for parametric testing. All data groups were determined to have equal variance as analyzed by Levene's test. Unless noted in Figure Legends, all data were assessed by One-Way ANOVA followed by a Tukey’s post test to determine significance.

#### Supplemental Figures

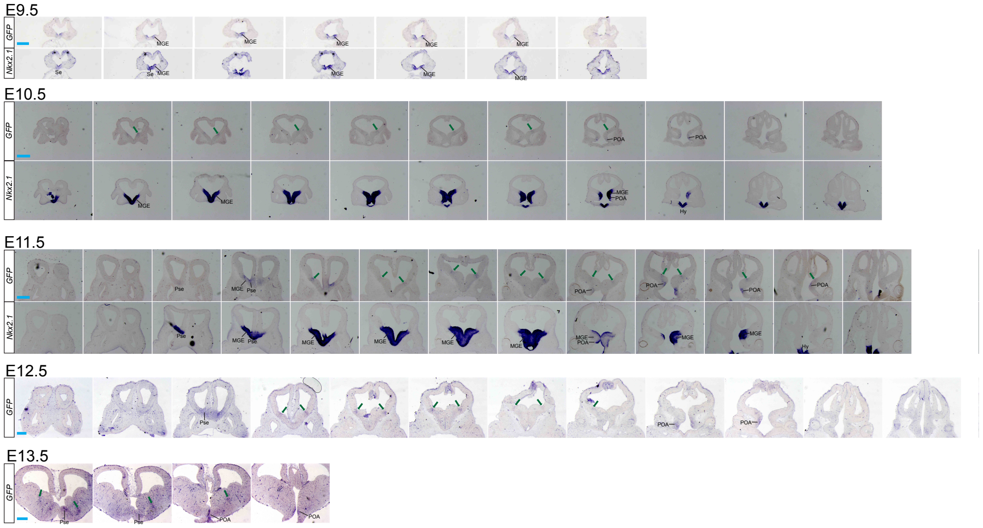


**Fig. S1. *1538* activity in the MGE/LGE sulcus, PSe, and POA.**
Data related to Figure 1. *Gfp in situ* hybridization (ISH) on E9.5 thru E13.5 coronal series reporting *1538* activity. *Nkx2.1* ISH on adjacent sections in E9.5 thru E11.5 coronal series. *Gfp* expression is in the MGE/LGE sulcus through E13.5 (green arrow; also known as MGE0/1 (*2*)). Hy, hypothalamus; MGE, medial ganglionic eminence; POA, preoptic area, PSe, paraseptal; Se, septum. Blue scale bar, 200 uM.

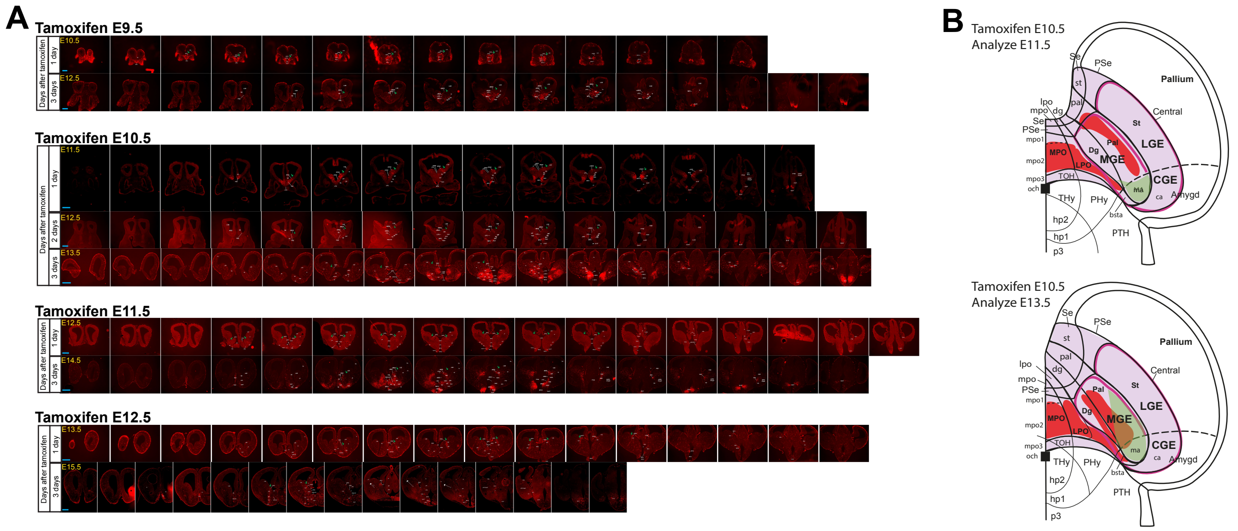


### Fig. S2. Fate mapping of *1538* lineages

Data related to Figure 1. **(A)** Coronal series showing fate maps of E9.5, E10.5, E11.5, and E12.5 *1538* lineages (labelled by tdTomato) one, two, and three days after tamoxifen treatment. Green arrow, MGE/LGE sulcus. Blue scale bar, 200 uM. **(B)** Schema representing VZ/SVZ location (red regions) of E10.5 *1538* lineage at E11.5 (top) and E13.5 (bottom). Tamoxifen treatment at E10.5. The topological schema of the forebrain is based on (*31*). Red regions, VZ/SVZ location of *1538* lineage. Lavender region, subpallium. Black solid lines, delineate subpallial subdivisions. Black dash lines, caudal domains contributing to the amygdala and CGE. Black square, optic chiasm. Purple lines, MGE/LGE/CGE domains. Green region, interneuron producing domain. Uppercase labels denote VZ/SVZ while lowercase labels denote MZ. Note the caudoventral movement of the tdTomato$^{+}$ region from the pallidal (Pal) toward the diagonal (Dg) domain. Also, note the tdTomato$^{+}$ preoptic regions (LPO and MPO) do not move except into bsta. Finally, note the rostral expansion of the interneuron producing domain (green). Abbreviations: ac, anterior commissure; acb, nucleus accumbens; A-HY, alar hypothalamus; B-HY, basal hypothalamus; bsta, bed nucleus stria terminalis amygdalar part; bstl, bed nucleus stria terminalis lateral; bstm, bed nucleus stria terminalis medial; bstp, bed nucleus stria terminalis posterior part; ca, central amygdala; cdMGE, caudodorsal MGE; CGE, caudal ganglionic eminence; ch. fiss, choroid fissure; cMGE, caudal MGE; db, diagonal band; dbh, horizontal limb of the diagonal band; dg, diagonal region; cx, cortex; dMGE, dorsal medial ganglionic eminence; fi, fimbria of the hippocampus; gp, globus pallidus; gpe, globus pallidus externa; gpi, globus pallidus interna; hi, hippocampus; hp1, hypothalamic prosomere 1; hp2, hypothalamic prosomere 2; ic, internal capsule; in, interneurons; ivf, interventricular foramen; LGE, lateral ganglionic eminence; LPO, lateral preoptic area; lt, lamina terminalis; ma, medial amygdala; mbv, meningial blood vessels; men, meninges; MPO, medial preoptic area; och, optic chiasm; OP, optic vesicle; os, optic stalk; ot, optic tract; p3, prosomere 3; Pal, pallidal domain; pall amygd, pallial amygdala; ped, cerebral peduncle; PHy, peduncular hypothalamus; PSe, paraseptal domain; pth, prethalamus; PThE, prethalamic eminence; rMGE, rostral MGE; Se, septal domain; si-d, diagonal substantia innominata domain; si-p, pallidal substantia innominata domain; st, striatum; stria term, stria terminalis; THy, terminal hypothalamus; TOH, telencephalo-opto-hypothalamic transition zone; vMGE, ventral medial ganglionic eminence; TH, thalamus; vdb, vertical limb of the diagonal band; vpal, ventral pallidal domain; vpal-p, pallidal ventral pallidal domain; vpal-pse, paraseptal ventral pallidal domain; vpal-se, septal ventral pallidal domain; zli, zona limitans.

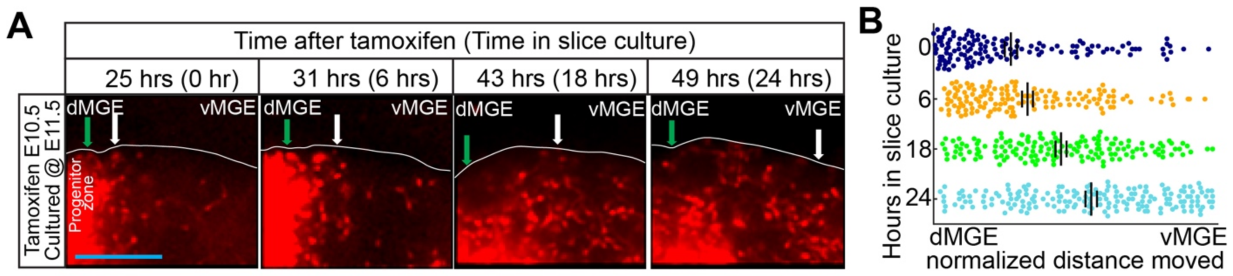


**Fig. S3. MGE progenitors in the *1538* lineage move ventrally in the VZ in slice cultures**
Data related to Figure 1. **(A)** *1538* lineage labelled with tdTomato$^+$ (tamoxifen at E10.5) in coronal hemisections from E11.5 MGE. The sectioned MGE was placed in culture for 0, 6, 18, and 24 hours. Time 0 hours is one hour after E11.5 brain slices were cultured. Green arrow, MGE/LGE sulcus. White arrow, approximate center of tdTomato$^+$ cells in the VZ moving ventrally from the MGE/LGE sulcus. Blue scale bar, 100 uM. **(B)** Measurement of tdTomato$^+$ cell (*1538* lineage) movement along the progenitor zone (VZ/SVZ) 0, 6, 18, and 24 hours in culture. Individual tdTomato$^+$ cells in the progenitor zone were plotted from the MGE/LGE sulcus (green arrow) to the MGE/POA sulcus. The distance between the MGE/LGE sulcus and MGE/POA sulcus was normalized. Experiment was performed on 3 separate brains (N=3); about 180 to 220 cells were analyzed per time point. Error bars, SEM. Abbreviations: dMGE, dorsal medial ganglionic eminence; vMGE, ventral medial ganglionic eminence.

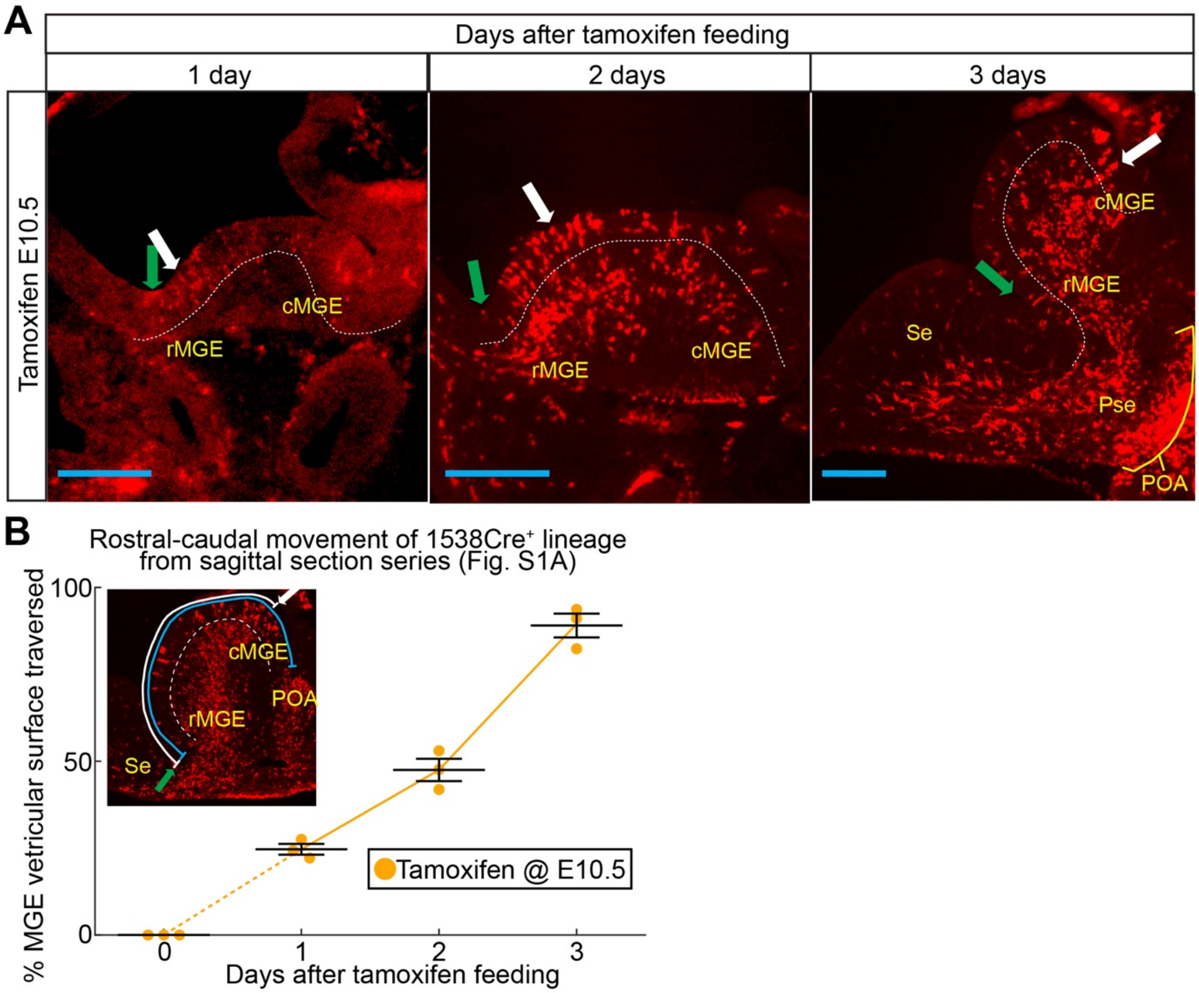


**Fig. S4. MGE progenitors in the *1538* lineage move caudally in the VZ**

Data related to Figure 1. **(A)** Fate mapping of *1538* lineages (tdTomato$^+$; tamoxifen given at E10.5) one, two, and three days after tamoxifen treatment. tdTomato$^+$ progenitors (white arrow) move caudally away from the MGE/LGE sulcus (green arrow) along the MGE VZ. Blue scale bar, 100 uM. **(B)** *1538* progenitor movement along the VZ is measured one, two, and three days after tamoxifen treatment (orange line) (n=3 for each time point). Inset: tdTomato$^+$ cell movement along the VZ was calculated by measuring the distance (white line) between the approximate center of the 1538 progenitors (white arrow) and the MGE/Septal sulcus (green arrow). The distance of the tdTomato$^+$ cell movement was normalized to the distance between the Septal/MGE and MGE/POA sulcus (blue line). Error bars, SEM. White dash line, MGE SVZ and MZ boundary. Abbreviations: cMGE, caudal medial ganglionic eminence; POA, preoptic area; PSe, paraseptal domain; rMGE, rostral medial ganglionic eminence; Se, septal domain.

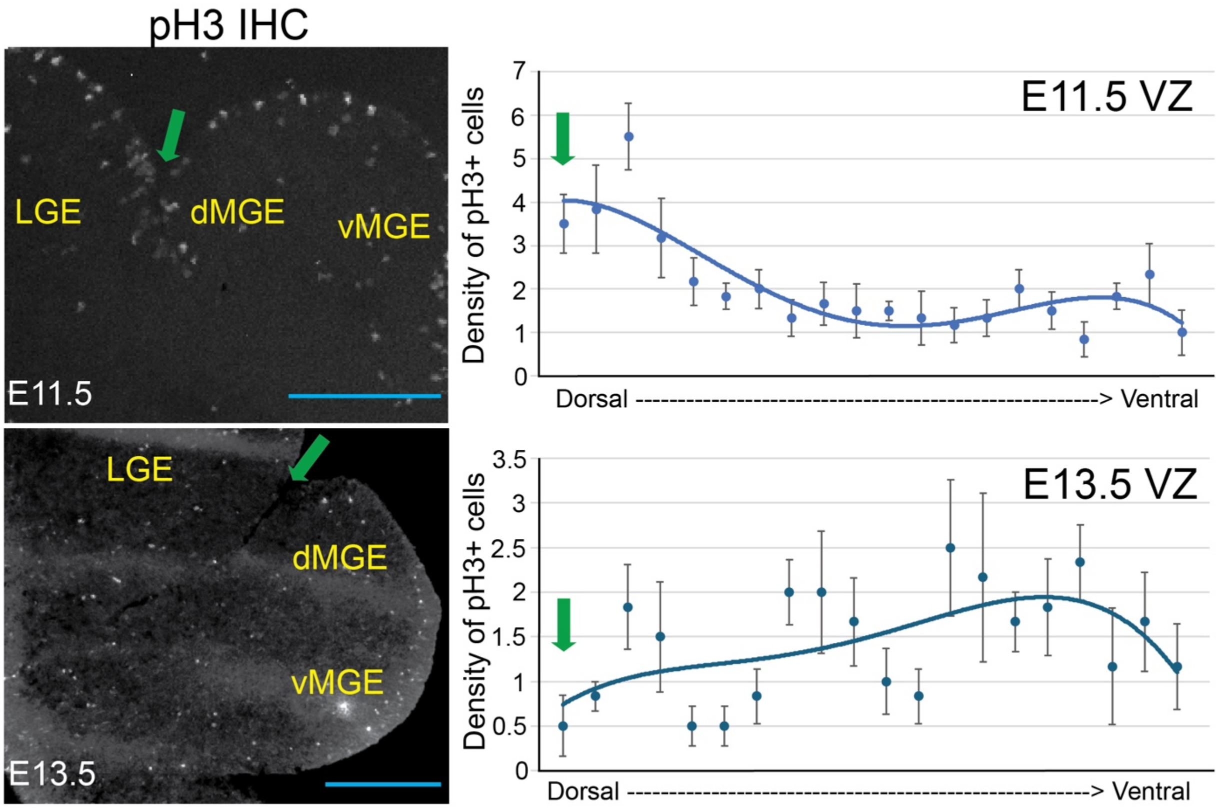


**Fig. S5. MGE/LGE Sulcus (MGE0/1) has the highest density of pH3⁺ cells in the E11.5 MGE**

Data related to Figure 2. Phospho-histone H3 (pH3) immunohistochemistry on E11.5 (upper left panel) and E13.5 (lower left panel) coronal wildtype sections. Blue scale bar, 200 uM. pH3 density measurement along the E11.5 (upper right panel) and E13.5 (lower right panel) MGE VZ (n=6 for each time point). Density was counted in 20 equally spaced bins along the dorsal-ventral MGE VZ axis. Green arrow: MGE/LGE sulcus (MGE0/1). Note, higher pH3 density in the MGE/LGE sulcus at E11.5 but not at E13.5. Error bars, SEM. Abbreviations: LGE, lateral medial ganglionic eminence; dMGE, dorsal medial ganglionic eminence; vMGE, ventral medial ganglionic eminence; VZ: ventricular zone.

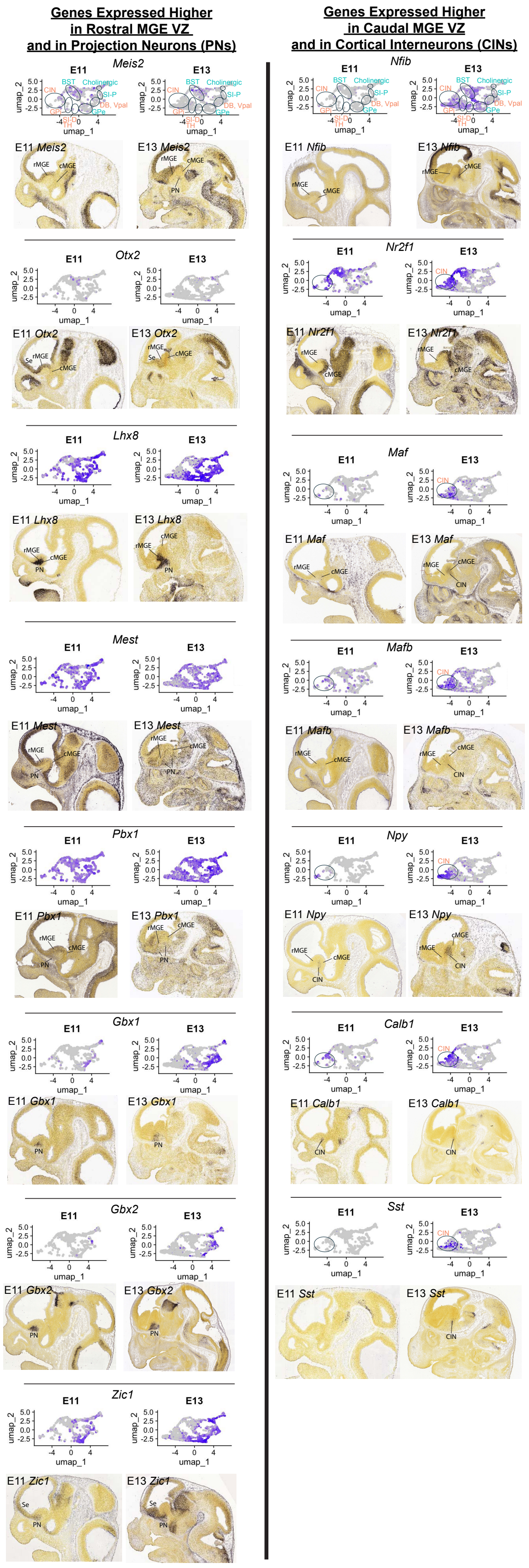

Genes Expressed Higher in Rostral MGE VZ and in Projection Neurons (PNs)
Genes Expressed Higher in Caudal MGE VZ and in Cortical Interneurons (CINs)
Meis2
Otx2
Lhx8
Mest
Pbx1
Gbx1
Gbx2
Zic1
Nfib
Nr2f1
Maf
Mafb
Npy
Calb1
Sst
E11
E13
umap_1
umap_2
CIN
BST
Cholinergic
SI-P
DB, Vpal
SPe
E11 Meis2
E13 Meis2
E11 Otx2
E13 Otx2
E11 Lhx8
E13 Lhx8
E11 Mest
E13 Mest
E11 Pbx1
E13 Pbx1
E11 Gbx1
E13 Gbx1
E11 Gbx2
E13 Gbx2
E11 Zic1
E13 Zic1
E11 Nfib
E13 Nfib
E11 Nr2f1
E13 Nr2f1
E11 Maf
E13 Maf
E11 Mafb
E13 Mafb
E11 Npy
E13 Npy
E11 Calb1
E13 Calb1
E11 Sst
E13 Sst
rMGE
cMGE
PN
Se

**Fig. S6. Correspondence between regional gene expression in the MGE VZ, and the types of neurons produced**

Data related to Figures 3 and 4. Gene expression for 15 genes that are preferentially expressed in MGE projection neurons (PN) (and rostral MGE VZ), or cortical interneurons (CINs) (and caudal MGE VZ) is shown at E11.5 and E13.5 using Feature Plots and ISH. Information about the feature plots of E11.5 and E13.5 1538 MZ cells can be found in Figures 4D and S7A. ISH is from the Allen Developing Brain Atlas; it shows expression at E11.5 and E13.5 sagittal sections showing genes highly expressed in rostral MGE VZ and in projection neurons (left side) or genes highly expressed in caudal MGE VZ and in cortical interneurons (right side). Blue labels in feature plots on the top row indicate cell types that are largely produced at E11.5, whereas orange labels indicate cell types that are largely produced at E13.5. Abbreviations: bst, bed nucleus stria terminalis; cMGE, caudal MGE; db, diagonal band; CIN, cortical interneuron; gpe, globus pallidus externa; gpi, globus pallidus interna; PN, projection neuron; rMGE, rostral MGE; Se, septum; si-d, diagonal regions of the substantia innominata; si-p, pallidal region of the substantia innominata; TH, tyrosine hydroxylase$^{+}$ neurons; vpal, ventral pallidum.

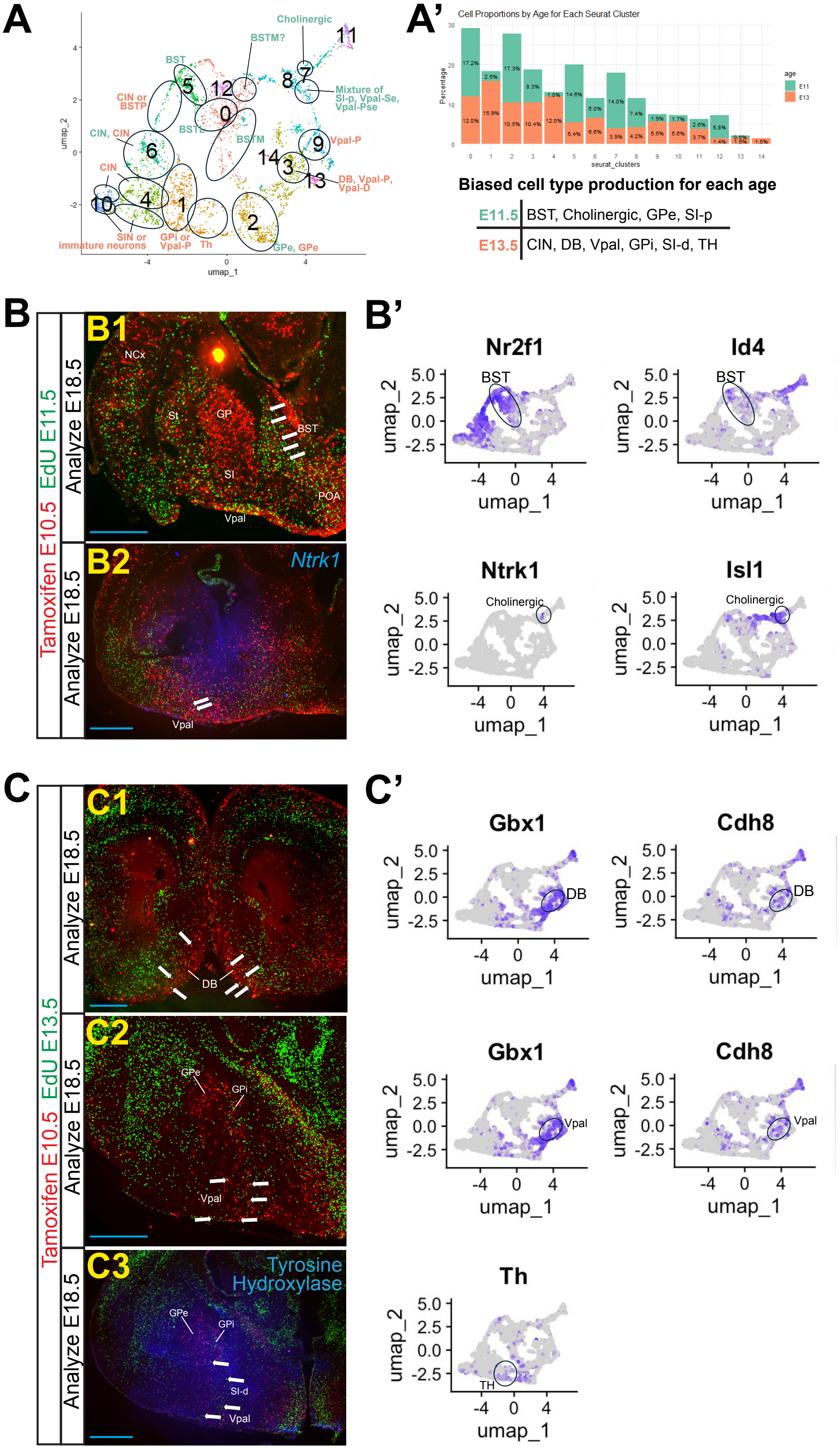
A
Cholinergic
BSTM?
BST
CIN or BSTP
CIN, CIN
CIN
BSTL
BSTM
Mixture of SI-p, Vpal-Se, Vpal-Pse
Vpal-P
DB, Vpal-P, Vpal-D
SIN or immature neurons
GPi or Vpal-P
Th
GPe, GPe
umap_1
umap_2
A'
Cell Proportions by Age for Each Seurat Cluster
Percentage
seurat_clusters
age
E11
E13
Biased cell type production for each age
E11.5 BST, Cholinergic, GPe, SI-p
E13.5 CIN, DB, Vpal, GPi, SI-d, TH
B
Tamoxifen E10.5 EdU E11.5
B1
Analyze E18.5
NCx
St
GP
BST
SI
POA
Vpal
B2
Ntrk1
Analyze E18.5
Vpal
B'
Nr2f1
Id4
Ntrk1
Isl1
BST
Cholinergic
C
Tamoxifen E10.5 EdU E13.5
C1
Analyze E18.5
DB
C2
Analyze E18.5
GPe
GPi
Vpal
C3
Tyrosine Hydroxylase
Analyze E18.5
GPe
GPi
SI-d
Vpal
C'
Gbx1
Cdh8
DB
Vpal
Th
TH

**Fig. S7. Progenitors in the *1538* lineage produce neurons of different cell fates as the progenitors move caudoventrally along the VZ of the MGE**

Data related to Figure 4. **(A)** *Dcx*$^+$ MZ cells from the UMAP in Figure 3A were reclustered in this UMAP (left panel). Blue labels indicate cell types that are largely produced at E11.5 whereas orange labels indicate cell types that are largely produced at E13.5. Black ovals approximate the boundaries between cell types. Cell clusters 1-14 each have their own arbitrary color. Cell-type identity was assigned for each UMAP region based on expression markers (table S4). **(A')** Histogram showing proportion of E11.5 and E13.5 MZ cells for each cell cluster (right panel). Below are listed cell types preferentially produced at E11.5 vs E13.5. **(B)** Identification of MGE cells generated at E11.5. E18.5 merged coronal hemisections of EdU$^+$ (green) (E11.5 injection of EdU), and *1538* lineage labeled by tdTomato$^+$ IHC (red) (E10.5 tamoxifen). (B1) White arrows show EdU/tdTomato-co-expressing cells in the BST. (B2) Triple-labeling with *Ntrk1* ISH (blue). White arrows show triple-labeled cells in the Vpal. Blue scale bar, 200 uM. **(B')** Feature plots of E11.5 and E13.5 *1538*-lineage MZ cells showing the expression of markers for BST neurons (top, *Nr2f1* and *Id4*) or for Cholinergic neurons (bottom, *Ntrk1* and *Isl1*). **(C)** Identification of MGE cells generated at E13.5. E18.5 merged coronal hemisections of EdU$^+$ (green) (E13.5 injection of EdU), and *1538* lineage labeled by tdTomato$^+$ IHC (red) (E10.5 tamoxifen). (C1) White arrows show EdU/tdTomato-co-expressing cells in the DB. (C2) White arrows show EdU/tdTomato-co-expressing cells in Vpal. (C3). Triple-labeling for Tyrosine Hydroxylase (TH) IHC (blue). White arrows show triple-labeled cells in the Vpal and SI-d. Blue scale bar, 200 uM. **(C')** Feature plots of E11.5 and E13.5 *1538*-lineage MZ cells showing the expression of markers for DB and Vpal neurons (*Gbx1* and *Cdh8*) and TH$^+$ neurons (*Th*). Abbreviations: BST, bed nucleus stria terminalis; BSTL, bed nucleus stria terminalis lateral; BSTM, bed nucleus stria terminalis medial; BSTP, bed nucleus stria terminalis posterior; CIN, cortical interneurons; DB, diagonal band; GPe, globus pallidus externa; GPi, globus pallidus interna; Ncx, neocortex; POA, preoptic area; SI-d, diagonal domain of the substantia innominata; SI-p, pallidal domain of the substantia innominata; sin, striatal interneuron; St, striatum; Th, tyrosine hydroxylase$^+$ neurons; Vpal, ventral pallidum; Vpal-D, diagonal domain of the ventral pallidum; Vpal-P, pallidal domain of the ventral pallidum; Vpal-Pse, paraseptal domain of the ventral pallidum; Vpal-Se, septal domain of the ventral pallidum.

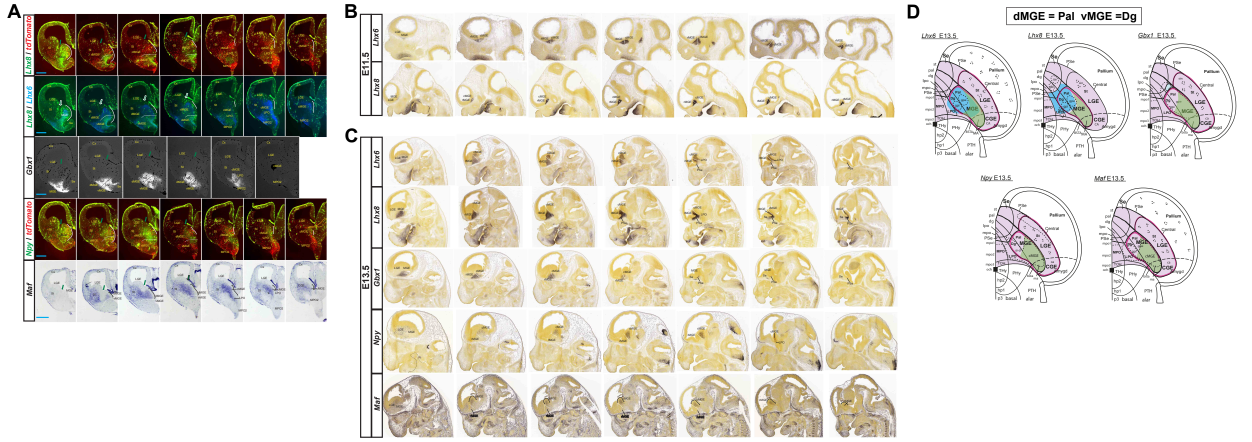


**Fig. S8. *Lhx8*-negative interneuron producing domain in the E13.5 MGE**

Data related to Figure 4. **(A)** E13.5 coronal series showing MGE expression of 6 genes that highlights the interneuron producing domain (IN domain; brackets in the *Maf* ISH in the fifth row). Row 1: *Lhx8* (green) fluorescent ISH co-stained with tdTomato (IHC; red, E10.5). tdTomato marks the 1538 lineage (Tamoxifen was given at E10.5). Red cells in the VZ are highlighted by a white line along the VZ surface. Row 2: *Lhx8* fluorescent ISH (green) with *Lhx6* fluorescent ISH (blue) (second row); Row 3: *Gbx1* fluorescent ISH; Row 4: *Npy* (green) fluorescent ISH with tdTomato (red, E10.5 1538 lineage); Row 5 *Maf* ISH (fifth row). Green arrow: MGE/LGE sulcus. Dashed line, dMGE and vMGE boundary. Blue scale bar, 100 uM. **(B)** ISH (Allen Developing Brain Atlas) showing *Lhx6* (first row) and *Lhx8* (second row) expression at E11.5 in sagittal sections. Projection neurons but not cortical interneurons are present in both the rostral and caudal MGE MZ based on *Lhx8*. Dashed line, rostral MGE (rMGE) and caudal MGE (cMGE) boundary. **(C)** Interneuron producing (IN) domain is in the caudal MGE (brackets in the *Maf* ISH in the fifth row). ISH (Allen Developing Brain Atlas) showing expression at E13.5 in sagittal sections of the same 5 genes shown in part A that are expressed in MGE SVZ/MZ. Projection neurons are present in both the rostral and caudal MGE MZ based on *Gbx1* (but not in the caudal MGE SVZ, where IN genes *Npy* and *Maf* expression are high). First row: *Lhx6*; Second row: *Lhx8*; Third row: *Gbx1*; Fourth row: *Npy*; and Fifth row: *Maf*. Dashed line, rostral MGE (rMGE) and caudal MGE (cMGE) boundary. **(D)** Schemas representing *Lhx6*, *Lhx8*, *Gbx1*, *Npy*, and *Maf* expression at E13.5. The topological schema of the forebrain is based on (*31*). Dashed ovals, denote MZ expression of each gene. Lavender region, subpallium. Black solid lines, delineate subpallial subdivisions. Purple lines, MGE/LGE/CGE domains. Uppercase labels denote VZ/SVZ regions while lowercase labels denote MZ regions. Blue region, VZ/SVZ expression of *Lhx6* and *Lhx8* (note that only *Lhx6* expression overlaps with the green region). Green region, interneuron producing domain. Purple outline, MGE/LGE/CGE domains. Black dash line, caudal domains contributing to the amygdala and CGE. Black square, optic chiasm. Abbreviations: al, ansa lenticularis; bsta, bed nucleus stria terminalis amygdalar part; ca, central amygdala; CGE, caudal ganglionic eminence; cMGE, caudal MGE; Cx: cortex; db, diagonal band; dg, diagonal region; dMGE, dorsal medial ganglionic eminence; gpe, globus pallidus externa; gpi, globus pallidus interna; hp1, hypothalamic prosomere 1; hp2, hypothalamic prosomere 2; ise, intermediate septum; LGE, lateral ganglionic eminence; LPO, lateral preoptic area; ma, medial amygdala; MPO2, medial preoptic area 2; MZ: mantle zone; och, optic chiasm; p3, prosomere 3; Pal, pallidal region; PHy, peduncular hypothalamus; PSe, paraseptal region;

pth, prethalamus; rMGE, rostral MGE; Se, septal region; si-d, diagonal region of substantia innominata; si-p, pallidal region of substantia innominata; sin, striatal interneuron; st, striatum; THy, terminal hypothalamus; TOH, telencephalo-opto-hypothalamic transition zone; vMGE, ventral medial ganglionic eminence.

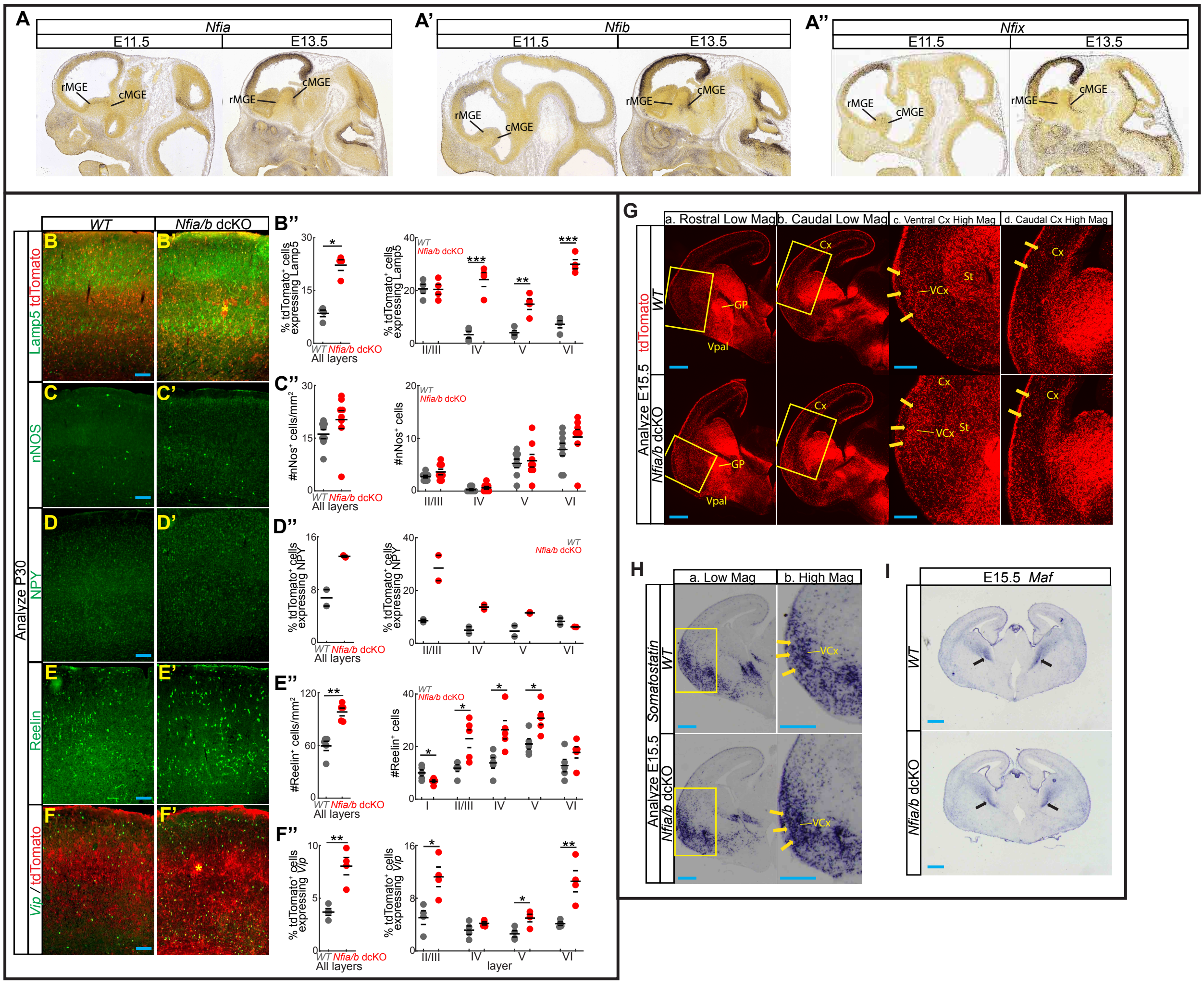


**Fig. S9.** ***Nfia* and *Nfib* temporally regulate the number and identity of CINs**
Related to Figure 5. **(A)** *Nfia* (A), *Nfib* (A'), and *Nfix* (A'') ISH at E11.5 and E13.5 on sagittal sections (from Allen Developing Brain Atlas). Note increase in *Nfia*, *Nfib*, and *Nfix* expression in cMGE at E13.5. cMGE, caudal MGE; rMGE, rostral MGE. **(B-F)** Lamp5 IHC (green) co-stained with tdTomato (red) (B, B'), nNos IHC (green) (C, C'), NPY IHC (green) (D, D'), Reelin IHC (green) (E, E'), and *Vip* ISH co-stained with tdTomato (red) (F, F') on P30 somatosensory neocortex from wildtype (B-F) and *Nfia/b* dcKO (B'-F') mice. CINs are red because of tdTomato labeling using *Nkx2.1-Cre.* Asterisk, accumulation of tdTomato$^{+}$ cells occasionally seen in the neocortex of *Nfia/b* dcKO mice. Blue scale bar, 100 uM. **(B'')** Percentage of tdTomato$^{+}$ cells expressing Lamp5. **(C'')** nNos$^{+}$ density. **(D'')** Percentage of tdTomato$^{+}$ cells expressing NPY. **(E'')** Reelin$^{+}$ density. **(F'')** Percentage of tdTomato$^{+}$ cells expressing *Vip* (n=4, 8, 2, 5, and 4, respectively, for each genotype). Quantifications for all layers are in the left panels. Quantifications for individual cortical layers are in the right panels. *, p<0.05; **, p<0.01, ***, p<0.001. Error bars, SEM. **(G)** tdTomato IHC on E15.5 rostral (Column a) and caudal (Column b) coronal hemisections from wildtype (top row) and *Nfia/b* dcKO (bottom row) mice. Yellow box in Column a indicates ventral cortical region magnified in column c. Yellow box in Column b indicates caudal cortical region magnified in column d. Arrows point to

increased densities of tdTomato$^{+}$ cells in the ventral (Column c) and caudal (Column d) cortices of *Nfia/b* dcKO mice. Blue scale bar, 200 uM. **(H)** *Somatostatin* ISH on E15.5 coronal hemisections (Column a) from wildtype (top row) and *Nfia/b* dcKO (bottom row) mice. Yellow box in Column a indicates ventral cortical region magnified in column b. Arrows point to increased densities of *Sst*$^{+}$ cells in the ventral cortex (Column b) of *Nfia/b* dcKO mice. Blue scale bar, 200 uM. **(I)** *Maf* ISH on E15.5 coronal sections from wildtype (F) and *Nfia/b* dcKO (I') mice. Black arrows, *Maf* expression in MGE SVZ. Blue scale bar, 200 uM. Cx, cortex; GP, globus pallidus; St, striatum; Vcx, ventral cortex; Vpal, ventral pallidum.

**Table S1. Top genes expressed in E11.5 and E13.5 1538 MGE VZ cells.** Top E11.5 VZ genes have positive average $Log_2$ fold change values. Top E13.5 VZ genes have negative average $Log_2$ fold change values. Hyperlinks to the Allen Developing Brain Atlas (ABA) for embryonic ISH images of corresponding genes are provided, if available.

**Table S2. Top genes expressed in E11.5 and E13.5 1538 MGE MZ cells.** Top E11.5 MZ genes have positive average $Log_2$ fold change values. Top E13.5 MZ genes have negative average $Log_2$ fold change values. Hyperlinks to the Allen Developing Brain Atlas (ABA) for embryonic ISH images of corresponding genes are provided, if available.

**Table S3. List of 52 regulators and markers of projection neuron and cortical interneuron development used to recluster 1538 MGE MZ cells.**

**Table S4. Combinatorial markers of specific cell types used to identify MGE MZ clusters.**
